# The structure of Egalitarian in complex with the *K10* mRNA localization signal reveals a modular binding surface required for function

**DOI:** 10.1101/2025.08.03.668247

**Authors:** Zebin Hong, Li Jin, Jonas Mühle, Fulvia Bono

**Author notes:** Co-corresponding authors: Zebin Hong and Fulvia Bono. Contact. Paul Scherrer Institute, OFLG/102, 5232 Villigen, CH. Institute of Molecular and Cell Biology (IMCB), Agency for Science, Technology and Research (A*STAR), 61 Biopolis Drive, Proteos, Singapore 138673, Republic of Singapore.

## Abstract

Asymmetric localization of mRNAs is a prevalent mechanism for spatial control of protein function and typically involves active transport by cytoskeletal motors. The mechanisms of recognition of localizing mRNAs by motor complexes are poorly understood. Egalitarian is an adaptor protein that binds localization signals in specific RNAs in *Drosophila* and recruits them to the dynein-dynactin complex for microtubule-based transport. We determined the crystal structure of Egalitarian in complex with the localization signal of the *K10* mRNA. Three structural units of Egalitarian, a 3’-5’exonuclease domain, a linker and a C-terminal domain form shape-specific, base-directed and backbone interactions with the RNA. Based on the structure we identified conserved residues recognizing RNA *in vitro*. Genome-edited flies with mutations in these residues have deficits in Egalitarian function that are congruent with changes in *in vitro* RNA binding affinities. Our work demonstrates how a minimal RNA localization signal is recognized by an RNA localization factor.

## Introduction

The localization of mRNAs to distinct cytoplasmic domains regulates many biological processes in eukaryotic cells such as embryonic patterning, cell polarity and neuronal function(Besse and Ephrussi, 2008; Holt and Bullock, 2009; Sutton and Schuman, 2006). This process is an effective way to restrict expression of proteins in cytoplasmic sub-compartments, thus spatially controlling gene expression(Martin and Ephrussi, 2009; St Johnston, 2005).

The transport of most localized mRNAs depends on cytoskeletal motors(Martin and Ephrussi, 2009; Tekotte and Davis, 2002). The localizing mRNAs are recognized by *trans*-acting protein factors that link them to microtubule-based motors. This recognition depends on *cis*-acting elements located mainly in the 3’ untranslated regions (3’UTRs) of mRNAs(St Johnston, 1995). The molecular mechanisms of how *trans*-acting factors recognize the RNA signals and link the mRNAs to the motor machinery are poorly understood.

Many mRNAs move towards the minus ends of microtubules by the dynein motor and its accessory complex, dynactin(Herbert et al., 2017; Trovisco et al., 2016; Vazquez-Pianzola et al., 2017). In *Drosophila melanogaster* the dynein-dynactin complex is involved in the localization of several essential patterning factors during early development. Dynein-dynactin-dependent transport has been described in several *Drosophila* cell types including oocytes, syncytial blastoderm embryos, neuroblasts and sensory neurons(Bullock and Ish-Horowicz, 2001; Ho et al., 2022; Hughes et al., 2004). mRNAs are linked to the dynein-dynactin machinery in these cells by the evolutionarily conserved coiled-coiled adaptor Bicaudal D (BicD) and the RNA-binding protein Egalitarian (Egl)(Bullock and Ish-Horowicz, 2001; Claussen and Suter, 2005; Dienstbier et al., 2009; Hughes et al., 2004; Mach and Lehmann, 1997; Navarro et al., 2004; Oh et al., 2000). Egl directly binds to mRNA localization elements in different mRNAs as well as the C-terminal domain (CTD) of BicD(Dienstbier et al., 2009). In addition, Egl also directly binds to the Dynein light chain (Dlc), facilitating cargo transport(Hoogenraad et al., 2001; Navarro et al., 2004) Transport complexes consisting of Egl, BicD, dynein, dynactin and mRNAs reconstituted *in vitro* are sufficient for long distance mRNA transport on microtubules(McClintock et al., 2018; Sladewski et al., 2018), revealing this is the minimal machinery for trafficking.

Minimal mRNA signals required for Egl-mediated minus-end-directed transport have been characterized in several mRNAs, including *bicoid (bcd)*, *fushi tarazu*, *gurken*, *hairy*, *fs(1)K10 (K10)*, *wingless*, and the *I Factor* retrotransposon(Bullock et al., 2003; Cohen et al., 2005, 2005; dos Santos et al., 2008; Macdonald and Kerr, 1998; Serano and Cohen, 1995; Snee et al., 2005; Van De Bor et al., 2005). These localization signals are predicted to form stem-loop structures of ∼40–65 nucleotides (nt)(Bullock et al., 2003; Cohen et al., 2005, 2005; dos Santos et al., 2008; Macdonald and Kerr, 1998; Serano and Cohen, 1995; Snee et al., 2005; Van De Bor et al., 2005). As these *cis*-acting elements show little primary sequence similarity to each other, it is unclear how they are recognized by the same protein.

*K10* is one of the most comprehensively studied mRNA cargoes for Egl. This mRNA is synthesized in the nurse cells and transported to the oocyte by Egl-BicD-dynein-dynactin where it is needed to establish dorso-ventral polarity of the future embryo(Cheung et al., 1992; Serano and Cohen, 1995). The 44-nucleotide transport and localization signal (TLS) of *K10* is necessary and sufficient for the localization of transcripts during early oogenesis (stage 2), late oogenesis (until stage 10) and in blastoderm embryos(Bullock and Ish-Horowicz, 2001; Serano and Cohen, 1995). Comprehensive mutagenesis of the *K10* TLS combined with injection-based localization assays in blastoderm embryos revealed that both primary and secondary structural elements are important for recognition by the transport machinery(Bullock et al., 2010).

The molecular details of how Egl recognizes the *K10* TLS or other RNA-localization elements are currently unknown. Egl is a 1004-amino-acid protein predicted to include a large uncharacterized N-terminal region, a 3’-5’ exonuclease (EXO) domain from the DEDDy superfamily and a C-terminal part containing a short consensus Dlc-binding region(Dienstbier et al., 2009; Mach and Lehmann, 1997; Navarro et al., 2004). Biochemical assays revealed that a large N-terminal portion of the protein encompassing the EXO domain is able to bind RNA(Dienstbier et al., 2009).

Here, we present the X-ray crystal structure of the minimal RNA-binding region of Egl in complex with the *K10* TLS. The structure reveals that RNA recognition is mediated by the EXO domain, a small positively charged domain and a helical linker separating these elements. Egl recognises the *K10* TLS by a combination of shape and sequence-specific interactions suggesting a mechanism for mRNA target discrimination. Editing of the *egl* locus in *Drosophila* using CRISPR/Cas9 shows that the residues involved in RNA recognition are essential for *in vivo* function, thus validating the structure.

## Results

### The central region of Egl is sufficient for RNA binding *in vitro*

Based on sequence analysis, three main conserved regions can be identified in Egl (**Figure 1A**; **Figure S1A**). The N-terminal region (Egl^N^, residues 1-500) includes an uncharacterized domain that bears the minimal binding region for BicD (BicD-BR, residues 1-79; (Dienstbier et al., 2009)), which links the RNP complex to dynein-dynactin(Dienstbier et al., 2009; Liu et al., 2013; Mach and Lehmann, 1997; Navarro et al., 2004). The central region of Egl (Egl^MID^, residues 500-850) is predicted to include a 3’-to-5’ exonuclease domain (EXO) that is essential for RNA-binding *in vitro* (Dienstbier et al., 2009), followed by a small domain predicted to contain helical features. The C-terminal portion of Egl (Egl^C^; residues 850-1004) is uncharacterized apart from and a short Dynein light chain-binding motif (Dlc-BM)(Hoogenraad et al., 2001; Navarro et al., 2004). Previous analysis of RNA binding *in vitro* showed that a protein encompassing Egl^N^ and Egl^Mid^ (Egl^1-819^) is sufficient for specific recognition of mRNA signals *in vitro*(Dienstbier et al., 2009).

**Figure 1.**
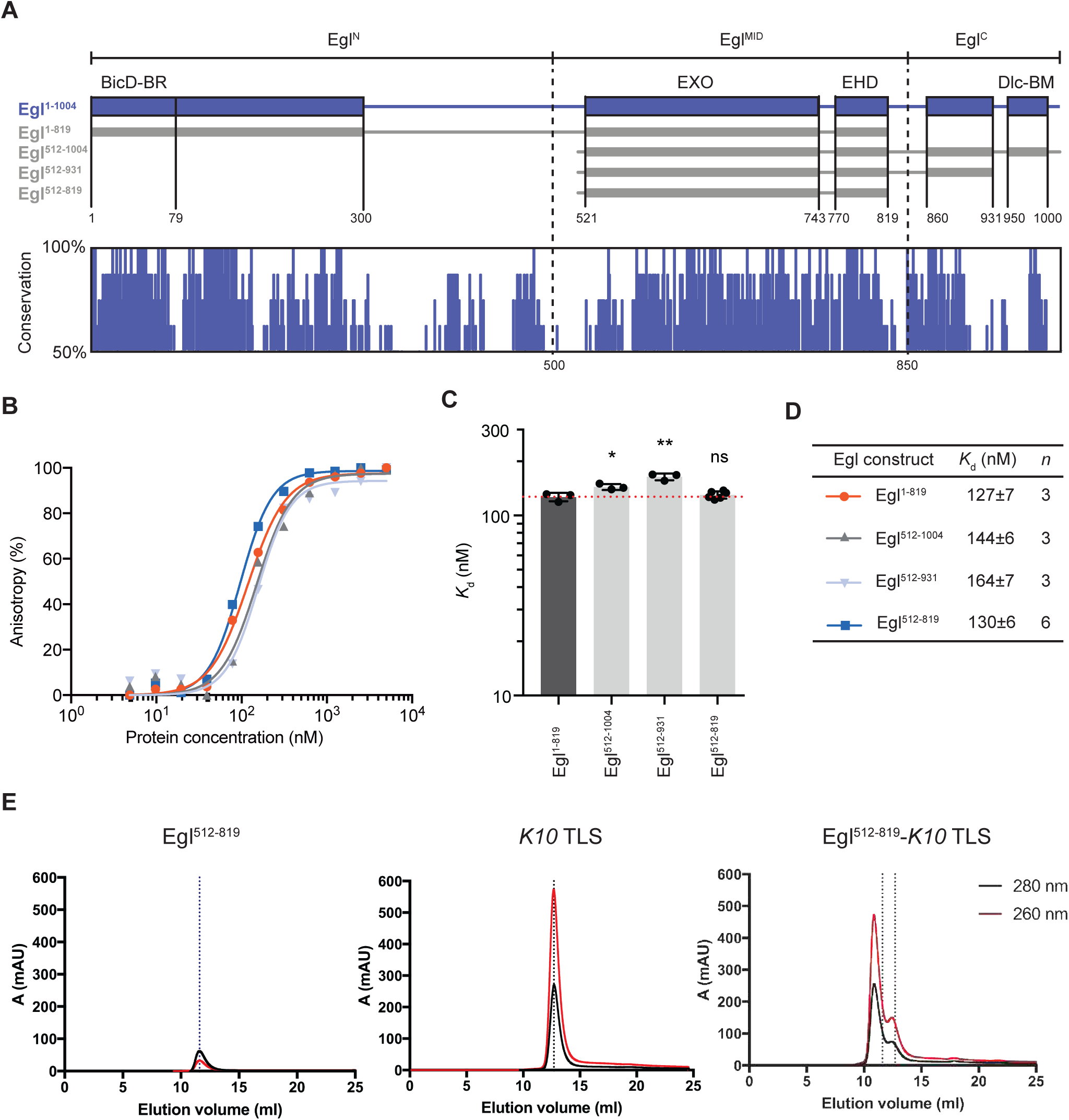
Egl^512-819^ is sufficient for RNA binding. **(A)** Schematic representation of the architecture of full-length Egl and the truncated constructs used. Solid boxes represent domains predicted to be folded, while lines are regions predicted to be unstructured. A conservation plot based on the alignment of multiple Egl homologs is shown below (Scores based on Consurf(Ashkenazy et al., 2016)). Sequences used to generate the alignment plot are shown in **Figure S1**. In the plot, residues positions are represented in the x-axis and percent identities (50-100%) are indicated in the y-axis. **(B)** *K*_d_ values were determined by FA. 5′-FAM-labeled *K10* TLS was incubated with increasing concentrations of Egl constructs and the data were fitted to the Hill equation to obtain the *K*_d_. **(C)** Column graph with mean *K*_d_ values shown as bars, standard deviation as black error bars and individual measurements as black dots. Data are represented as mean ± standard deviation (SD). Statistical significance was assessed using a two-sided unpaired Student’s t-test. ns, not significant; \**P* < 0.05, \*\**P* < 0.01, and \*\*\**P* < 0.001. ND, not determined. **(D)** Table of the *K10* TLS RNA binding affinities to various Egl truncations. *n* is the number of independent measurements. **(E)** SEC profiles of purified Egl^512-819^ (left), *K10* TLS alone (middle) and Egl^512-819^ preincubated with a 1.2-fold excess of *K10* TLS (right). The positions of the peaks are indicated by dotted lines in different shades of gray: the elution volume of Egl^512-819^ in isolation is in blue, while that of *K10* TLS RNA is in black

To further define the boundaries of the Egl RNA-binding region, we produced a series of truncations (**Figure 1A**), and measured binding affinities to the *K10* TLS by Fluorescence Anisotropy (FA). After confirming that the presence of a GST tag does not affect binding of Egl^512-819^ to RNA significantly in control experiments (**Figure S1B**), each Egl variant was purified as a GST fusion to improve solubility and enhance FA sensitivity. All truncated versions of Egl bind to *K10* TLS in the sub-micromolar range, similar to Egl^1-819^ (**Figure 1B-D; Table S1**), revealing that Egl^512-819^ is sufficient for full RNA binding. Including regions of the C-terminus (Egl^512-1004^) did not significantly affect RNA binding (**Figure 1B-D; Table S1**). Thus, we conclude that Egl^512-819^ is sufficient to bind to *K10* TLS *in vitro*.

### The Egl^512-819^ -*K10* TLS complex is stable and monomeric in solution

To understand the molecular mechanisms of Egl recognition of *K10* TLS, we first reconstituted a minimal complex *in vitro* using Egl^512-819^. We added two G-C-pairs at the base of the *K10* TLS stem that have been shown to stabilize the structure in solution and showed no loss of *in vivo* localization (Bullock et al., 2010). The RNA-protein complex is stable as shown by size-exclusion chromatography (SEC) and elutes as a well-defined peak with a larger elution volume compared with that of apo Egl^512-819^ (**Figure 1E**). Both Egl^512-819^ alone and in complex with *K10* TLS eluted with an elution volume consistent with that of a monomer. SEC coupled to right-angle light scattering (SEC-RALS) further showed that one Egl^512–819^ RNA-binding domain binds one *K10* TLS RNA element, establishing a 1:1 protein: RNA element stoichiometry for the minimal complex (**Figure S1E-F**). This result is consistent with the subsequent observation that individual RNA-binding sites within the dimeric Egl-BicD complex each engage a single structured RNA element (Singh et al., 2026).

### Overview of the structure of Egl^512-819^ in complex with *K10* TLS

We obtained crystals of the SEC-purified complex of Egl^512-819^-*K10* TLS. The structure was determined by molecular replacement (MR) at 3.1-Å resolution (**Table 1**) using a model generated by I-TASSER based on a homologous template, RNAse D(Barbosa et al., 2014). The crystal lattice includes two complexes of Egl^512-819^-*K10* TLS in the asymmetric unit (ASU). The two complexes superimpose with root mean square deviation (RMSD) of 0.634 Å over 1971 atoms. The crystal contact interface between them is stabilized through hydrogen bonds and salt bridges (**Figure S2A-C**).

**Table 1.** Data collection and refinement statistics of the crystal structures of Egl^512-819^-*K10* TLS complex and of MBP-(GSAM)-L-EHD and of *K10*-TR.

| **Data collection** |  |  |  |
| --- | --- | --- | --- |
| Crystal | Egl<sup>512-819</sup>-K10 TLS | MBP-Egl<sup>743-819</sup> | K10-TR |
| Wavelength (Å) | 1.0000 | 1.0000 | 0.9999 |
| Space group | P212121 | P212121 | P41212 |
| $a, b, c$ (Å) | 68.90, 107.40, 140.80 | 52.80, 82.00, 120.20 | 70.20, 70.20, 81.30 |
| $\alpha, \beta, \gamma$ (°) | 90, 90, 90 | 90, 90, 90 | 90, 90, 90 |
| Resolution (Å) | 50-3.08 (3.27-3.08) | 50-2.20 (2.34-2.20) | 50-3.60 (3.82-3.60) |
| $R_{\text{meas}}$ | 0.17 (1.41) | 0.12 (1.37) | 0.31 (1.29) |
| $I/\sigma I$ | 8.07 (1.55) | 8.10 (1.01) | 5.29 (1.57) |
| Completeness (%) | 99.8 (99.1) | 99.2 (98.7) | 99.7 (98.9) |
| Redundancy | 7.02 (7.10) | 3.08 (3.08) | 6.21 (6.24) |
| No. unique reflections | 36673 (5931) | 50894 (8163) | 4519 (731) |
| CC(1/2) | 0.999 (0.702) | 0.996 (0.354) | 0.992 (0.928) |
| **Refinement** |  |  |  |
| R-factor | 0.2402 | 0.1933 | 0.2914 |
| R-free | 0.2916 | 0.2059 | 0.2990 |
| Average B value (Å<sup>2</sup>) | 117.00 | 54.77 | 79.31 |
| No. atoms |  |  |  |
| Protein | 5828 | 3330 | - |
| RNA | 1642 | - | 874 |
| Ligand/ion | 8 | 23 | - |
| Water | - | - | - |
| R.m.s. deviations |  |  |  |
| Bond lengths (Å) | 0.014 | 0.016 | 0.004 |
| Bond angles (°) | 2.026 | 2.028 | 0.860 |
| **Ramachandran plot (MolProbity)** |  |  |  |
| Favored (%) | 92.99 | 97.64 | - |
| Allowed (%) | 7.01 | 2.36 | - |
| Outliers (%) | 0 | 00 | - |

Statistics for the highest-resolution shell are shown in parentheses. Molprobity was used for validation.

In the final Egl^512-819^ model, the C-terminal region of the construct is not visible (aa 769-819) and several loop regions (aa 512-519, 552-553, 652-655, 720-723) are disordered and could not be modeled (**Figure 2A-B**). In the *K10* RNA model, the loop from A20 to C27 (loop^R^) is not well defined and was also not modeled. The C-terminal part of the protein contains a region that we termed the Egl helical domain (EHD). To further study the EHD that could not be modeled unambiguously, we generated a construct encoding residues 743 to 819 of Egl (Linker (L)-EHD) fused to the Maltose Binding Protein (MBP) as a crystallization driver (**Figure 2A**). We crystallized and determined the crystal structure of MBP-L-EHD at 2.5-Å resolution by MR using MBP as a search model (**Table 1**). The obtained electron density is well defined and allowed us to model residues 743 to 806 with confidence (**Figure 2C; Figure S2D**). The final Egl^512–819^–K10 TLS model was obtained by superimposing the two crystal structures using the shared rigid helical linker (L) as the alignment reference (**Figure 2E**).

**Figure 2.**
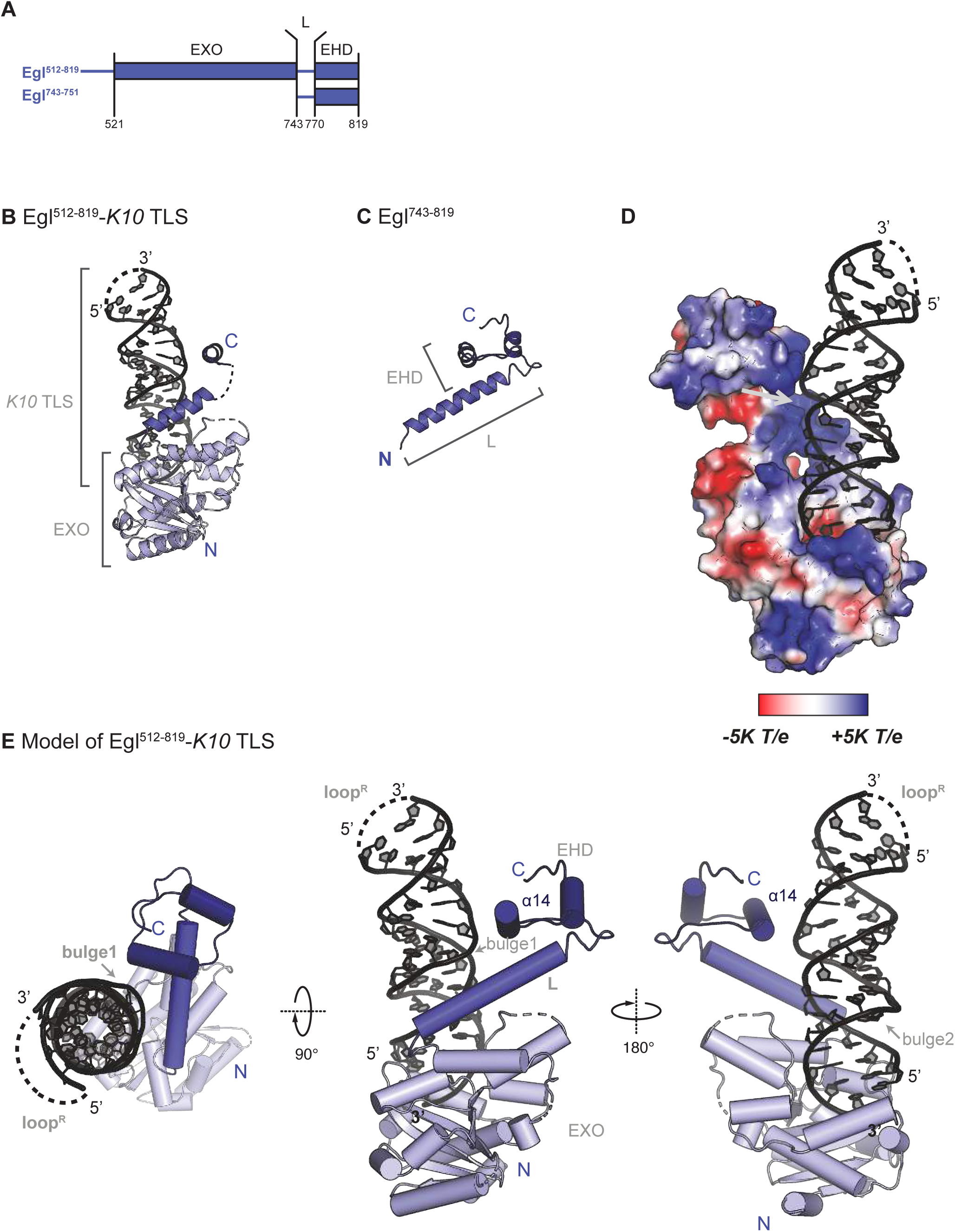
Structure of the Egl^512-819^-*K10* TLS complex. **(A)** Schematic representation of the Egl truncated constructs using for structural studies. **(B)** Crystal structure of Egl^512-819^ (with incomplete L and EHD regions) in complex with *K10* TLS. **(C**) Crystal structure of L-EHD. **(D)** Electrostatic potential plotted on the surface of Egl^512-81**9**^ with a gradient from red (negative) to blue (positive) from – to +5K *T/e*, respectively. The positively charged linker/EHD surface adjacent to K10 TLS Bulge1 was indicated with a gray arrow. **(E)** Model of Egl^512-819^ in complex with *K10* TLS in three different views related by 90° and 180° rotation. The model is obtained by superposition of the structures in **(B)** and **(C)**. The structures are shown as cartoon with Egl and *K10* TLS colored in a gradient in blue and black, respectively.

The overall structure of Egl^512-819^ consists of an N-terminal EXO domain and the small, positively charged EHD connected by a helical linker (**Figure 2E; Figure S2D**). *K10* TLS forms a typical A-form dsRNA duplex with an overall conformation similar to ideal A-form dsRNA (RMSD 3.1 Å over 40 nucleotides) (**Figure S3; Table1**). In the complex, RNA base pairing is mostly by Watson-Crick interactions with the exception of a wobble base pair (between G6-U43) and two bulges at C35 and A41 named Bulge1 and 2, respectively (**Figure 3B**). In the structure, Bulge2 lies on the opposite face of the helix to Bulge1, thus pointing to solvent and not interacting directly with the protein. Therefore, when in complex with Egl^512-819^, Bulge2 of *K10* TLS is in a similar structural position compared with *in silico* predictions (mFold;(Zuker, 2003)). However, it differs from the conformation of unbound *K10* TLS in solution, where Bulge2 is spatially arranged on the same side of the RNA stem as Bulge1 and involves nucleotide A39 (**Figure S3;**(Bullock et al., 2010)). Conversely, in the complex, Bulge2 consists of A41. Conformational flexibility could explain the less kinked conformation of the stem loop in the complex as compared with the previous NMR structure of *K10* TLS(Bullock et al., 2010). The folding of *K10* TLS in the complex is similar to a partial crystal structure we obtained of unbound *K10 TLS* that shows A-form conformation (**Figure S3; Table 1).**

**Figure 3.**
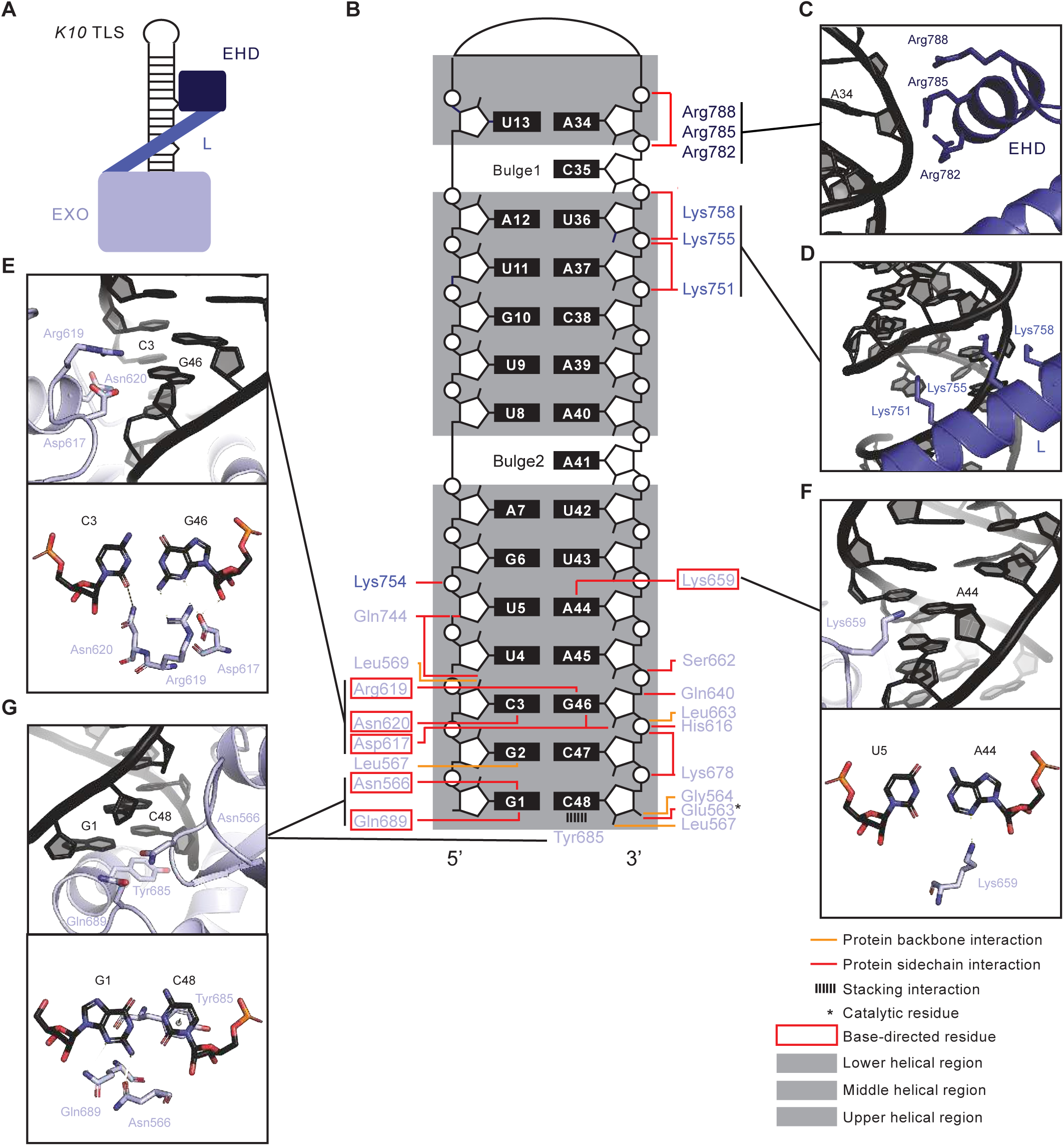
Protein-RNA interactions within the Egl^512-819^-*K10* TLS complex. **(A)** Schematic representation of the architecture of Egl^512-819^-*K10* TLS complex. Egl^512-819^ recognizes the *K10* TLS with several contacts in the EXO, Linker and EHD. **(B)** Schematic representation of the interactions between Egl^512-819^ and *K10* TLS. **(C-H)** The zoom-in panels show residues involved in the RNA binding. In **(C-H)**, Egl^512-819^ and *K10* TLS are shown as cartoon, and the figures in **(A-G)** share the same color-coding. Key hydrogen-bond and salt-bridge interactions are shown as dashed lines, with distances labelled in Å.

The interaction surface of Egl^512-819^ with *K10* TLS covers 905.7 Å^2^. Similarly, the *K10* TLS interaction surface involved in binding Egl^512-819^ is 1175.2 Å^2^. In the structure of the Egl^512-819^-*K10* TLS complex, the first 3’-most five nucleotides and the last 5’ complementary five at nt 44-48 associate tightly with the EXO domain (**Figure 2E and 3; Figure S2D**). The region that spans from nt C35 to A37, located around Bulge1, forms contacts with both linker and EHD. Interactions in this region are charged, and occur mostly with the RNA phosphate backbone (**Figure 2D and 3**).

The EXO domain adopts a typical α/β fold, consisting of a six-stranded β-sheet flanked by α-helices. A structure-based search using the DALI server(Holm and Laakso, 2016) identified several EXO domain-containing proteins including *E. coli* RNase D, *T. brucei* RRP6 and *B. mori* Exonuclease 3’-5’ domain-containing protein 1 (Exd1) (**Figure S4A**). Among these, RNase D shares the highest structural similarity with the EXO domain of Egl (RMSD 2.255 Å over 670 atoms) albeit with low sequence identity (21%) (**Figure S4A**). Based on structural comparisons, and as predicted by previous sequence analysis ((Navarro et al., 2004)Moser et al., 1997; **Figure S1**), the EXO domain of Egl belongs to the DEDDy superfamily. The conserved signature of DEDDy proteins are the catalytic tetrad and a tyrosine in so-called motif III. The EXO domain of Egl contains the four highly conserved acidic residues Asp561, Glu563, Asp621, and Asp708, and also the tyrosine (Tyr704) in motif III (**Figure S1**). The close arrangement of these residues in the Egl structure is consistent with the EXO domain of Egl retaining 3’-5’ exonuclease activity. However, mutation of all five putative catalytic residues does not affect RNA binding and *in vivo* function(Dienstbier et al., 2009; Navarro et al., 2004).

The EHD domain is composed of two positively charged α-helices facing each other in antiparallel arrangement and connected by unstructured loops (helix-loop-helix). Both helices contact the linker helix of Egl while helix α14 contacts the middle and upper regions of the RNA stem around bulge 1 (**Figure 2E and 3**). No significant structural homology of the EHD to known domains/motifs was retrieved upon databases searches but this could be due to its small size.

A 35 Å long helical linker connects the EXO and EHD domain, which is also highly positively charged. A comparison between Egl^512-819^-*K10* and the L-EHD structures shows that the linker is in a fixed position relative to the EXO and EHD domains and superposes with an RMSD of 1.031 Å over 117 atoms (**Figure 2)**. Indeed, the linker position is stabilized by several hydrophobic and salt bridges interactions with EHD and EXO domain residues (**Figure 2 and S4B**).

### Protein-RNA interactions within the Egl^512-819^-*K10* TLS complex

The association between Egl^512-819^ and the *K10* TLS RNA is mainly driven by electrostatic interactions. Positively charged residues are concentrated at the RNA interface (**Figure 2D**; **Figure S4C**). These positively charged residues are conserved in evolution (**Figure S4D**), supporting an important functional role.

The EXO domain clasps the lower part of *K10* TLS mainly by unspecific sugar- and phosphate-backbone interactions but also with a few direct base recognitions (**Figure 3**). The Watson-Crick base pairs G1-C48 and C3-G46 as well as A44 are involved in base-directed recognition. The side chains of Asn566 and Gln689 form hydrogen bonds with G1 and the aromatic side chain of Tyr685 is π-stacked against the 3’-terminal C48 (**Figure 3G; Figure S2D**). This interaction occurs with the first of the two RNA stabilizing G-C pairs we added to the sequence and involves a 3’OH base-specific recognition. However, the natural base pair predicted to form the base of the RNA stem is C-G, which would also be engaged in a similar and specific recognition in the absence of the artificial G-C doublet.

In addition, the conserved residues Asp617, Arg619 and Asn620 recognise the third base pair C3-G46 through hydrogen bonding interactions (**Figure 3E**). A direct interaction is also observed between Lys659 and A44 (**Figure 3F**). The nonspecific contacts between the Egl^512-819^ EXO domain and the RNA backbone can be divided into three groups: those forming hydrogen bonds with O2’ of the phosphate moiety (Gln744, Leu569 and Leu567), those forming hydrogen bonds or salt bridges with phosphate groups (Glu563, Lys678, Ser662, Leu663, His616, Gly564 and Glu563), and hydrophobic interactions with the sugar backbone (Gln640 and Gln744) (**Figure 3B**). Of the five putative catalytic residues only Glu563 interacts directly with the 3’-OH of the RNA backbone.

In the mid region of the RNA stem, four positively charged residues (Lys751, Lys754, Lys755 and Lys758) in the linker of Egl^512-819^ (**Figure 3D**) and three (Arg782, Arg785 and Arg788) in the EHD (**Figure 3C**) are in close contact with the negatively charged RNA backbone. Although there are no base-specific contacts in this region, the phosphate-backbone interaction in correspondence with Bulge1 at C35 is involved in electrostatic interactions with conserved positively charged amino acids and might play an important role in Egl substrate selectivity through a shape preference. There are no contacts between Egl^512-819^ and the upper region of *K10* TLS that include the upper stem and loop.

### Conserved residues of Egl^512-819^ are involved in RNA binding

The Egl EXO domain, linker and EHD domain all contact the *K10* TLS. To investigate the individual contribution of each region to RNA binding, we first generated a series of truncated versions of Egl that included the EXO domain (EXO, Egl^512-751^), EXO and linker (EXO-L, Egl^512-770^), and linker and EHD (L-EHD, Egl^743-819^) (**Figure 4A**) and measured binding affinities to 5’-6-carboxy-fluorescein (6-FAM)-labelled *K10* TLS by FA. The linker and EHD could not be purified separately and were not used for affinity measurements.

**Figure 4.**
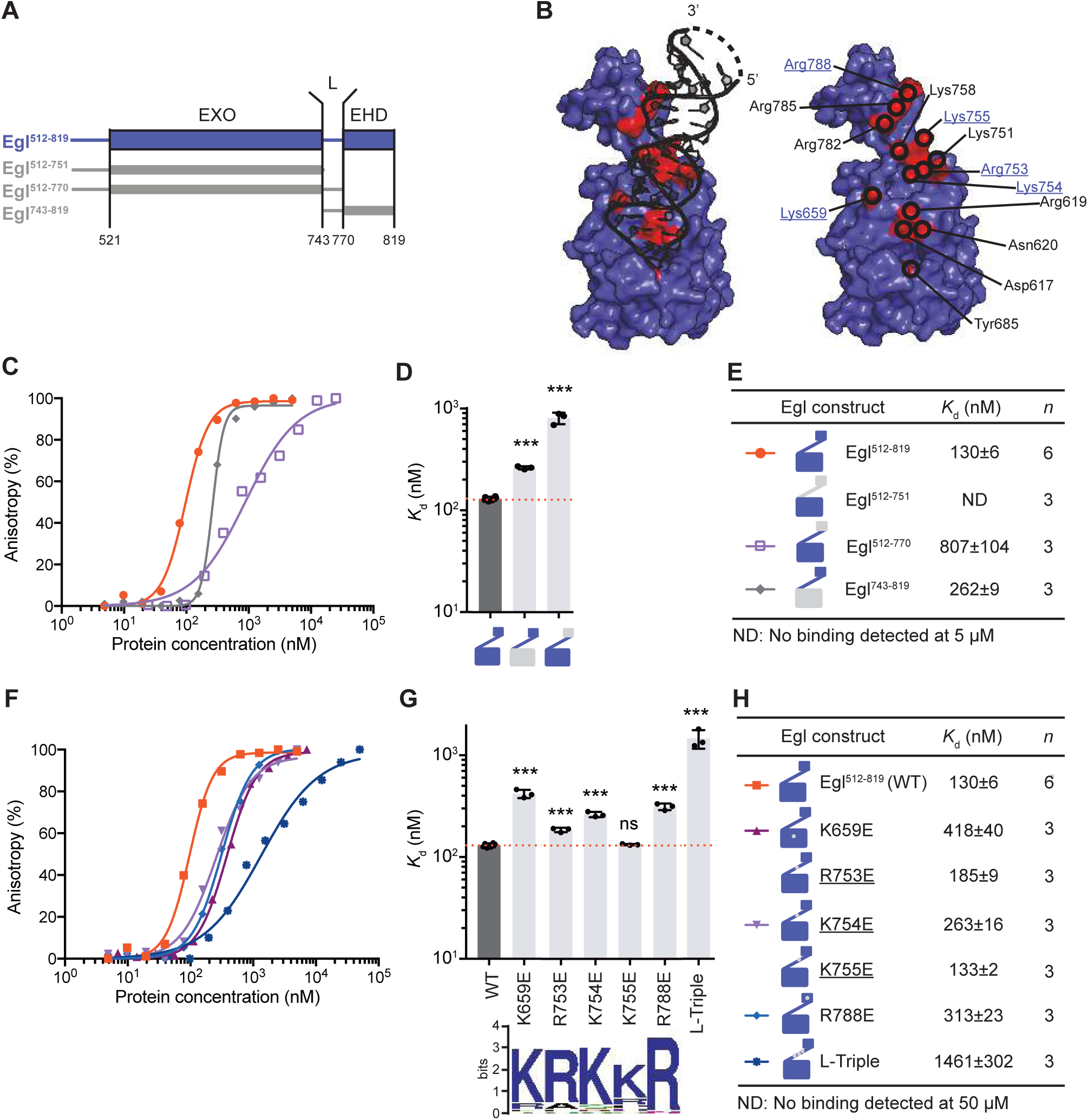
Mutagenesis analysis on the residues of Egl^512-819^ involved in RNA binding. **(A)** Schematic representation of the Egl truncated constructs. **(B)** The residues of Egl involved in the binding of *K10* TLS are highlighted in red and labeled on the protein surface rendering. On the right side, interacting residues are labelled. Underlined in blue are conserved residues mutated in the assays. **(C)** and **(F)** *K*_d_ values were determined by FA. 5′-FAM-labeled *K10* TLS was incubated with increasing concentrations of Egl mutants and the data were fitted to the Hill equation to obtain the *K*_d_. Data are represented as mean ± SD. **(D)** and **(G)** Column graph with mean *K*_d_ was shown as bars in the same color-coding, standard deviation as black error bars and individual *K*_d_ values as black dots. Weblogo representation of the residues involved in RNA binding was shown below the column graph. **(E)** and **(H)** *K*_d_ values of Egl mutants for *K10* TLS are shown in the table. Data are represented as mean ± SD with the number of independent measurements (n) indicated. Statistical significance was assessed using a two-sided unpaired Student’s t-test. ns, not significant; \**P* < 0.05, \*\**P* < 0.01, and \*\*\**P* < 0.001. ND, not determined.

The affinity of the Egl EXO domain for *K*10 TLS is at least 50 times lower than that of Egl^512-819^ as no binding up to 5 μM RNA could be detected. In contrast, L-EHD bound with a *K*_d_ of 262 ± 9 nM, only two-fold weaker than Egl^512-819^ (**Figure 4C-E; Table S1**). To examine the role of the linker region, we extended the EXO construct to include the linker. This EXO-L construct binds in the high nanomolar range with a *K*_d_ of 807 ± 104 nM, suggesting that the linker contributes to RNA binding and that the EHD acts as effector of RNA binding (**Figure 4C-E; Table S1**). Overall, these results indicate that Egl EXO domain, linker and EHD domain all contribute to binding, consistent with the structure.

To identify key residues involved in RNA-binding, we selected 16 conserved residues on each region for alanine scanning mutagenesis (**Figure 4B**). All mutants exhibit circular dichroism (CD) spectra similar to that of Egl^512-819^ wild type (wt), indicating that mutants are properly folded (**Figure S5A**). Most individual Ala mutations reduce the affinity to *K10* TLS only up to two-fold (**Figure S5B-C; Table S2**). Next, to disrupt salt bridges we individually replaced all of the positively charged residues by Glu. These reverse-charge mutants had either a similar or slightly stronger effect than the respective Ala mutants with an up to a 3-fold reduction in affinity (**Figure 4F-H; Table S2**). Next, we tried to combine single mutations to more strongly impair RNA binding. Three combinations (D617A/R619A/N620A, R753E/K754E/K755E and K659E/K754E/R788E) were selected for measurements. Asp617, Arg619 and Asn620 form base-directed hydrogen bonds with the base pair C3-G46 (**Figure 3A**). The D617A/R619A/N620A triple mutant shows two-fold reduction in binding with a *K*_d_ of 273 ± 12 nM (**Figure S5B-C; Table S2**). These residues are all located on the EXO domain and their mutation reduces binding affinity to a similar degree as the EXO-domain deletion. This result confirms the contribution of the EXO domain to RNA binding and reveals residues that functionally contribute to RNA binding. The next triple mutant we tested disrupts a positively charged patch (Arg753, Lys754 and Lys755) in the linker region (L-Triple). This mutant shows a ten-fold reduction in binding affinity (1461 ± 302 nM) (**Figure 4F-H; Table S2**). Finally, we combined mutations affecting each domain (EXO, K659E; linker, K745E; EHD, R788E). We did not detect RNA binding for this triple mutant (**Figure S5B-C; Table S2**).

### Determinants of RNA shape and sequence for Egl binding

To test the specificity of Egl^512-819^ binding towards RNA targets, we first measured direct binding affinity to different FAM-labeled RNA oligomers by FA. However, with this method, we could not detect large differences in affinities between Egl^512-819^ and various target and non-target RNAs (**Figure S6A-C; Table S3**). As an alternative approach, we set up a competition assay in which the preformed Egl^512-819^-FAM *K10* complex was incubated with increasing amounts of various unlabeled RNAs.

Previous work suggested that RNA secondary structure is important for specific recognition by Egl(Bullock et al., 2003; Bullock and Ish-Horowicz, 2001; Claussen and Suter, 2005; Hughes et al., 2004; Mach and Lehmann, 1997; Navarro et al., 2004; Oh et al., 2000). To test this, we measured the half-maximal inhibitory concentration (IC_50_) against single-stranded RNA (ssRNA), double-stranded RNA (dsRNA) and a non-target stem loop RNA derived from a bacteriophage (*MS2*) (**Figure 5A-C; S6D; Table S4**). ssRNA was a weak competitor of *K10* TLS for Egl^512-819^ binding with an IC_50_ of 112 ± 19 μM, which is 2000-fold weaker than control competitor *K10* TLS. In contrast, dsRNA and *MS2* stem loop are two-fold weaker competitors than *K10* TLS (**Figure 5A-C; Table S4**). Thus, a double helix is required for efficient competition with *K10* TLS. We also tested both ssDNA and dsDNA. As expected, dsDNA was a much weaker competition compared to dsRNA. ssDNA competition could not be detected even at a 40,000-fold excess over *K10* TLS (**Figure 5A-C; Table S4**).

**Figure 5.**
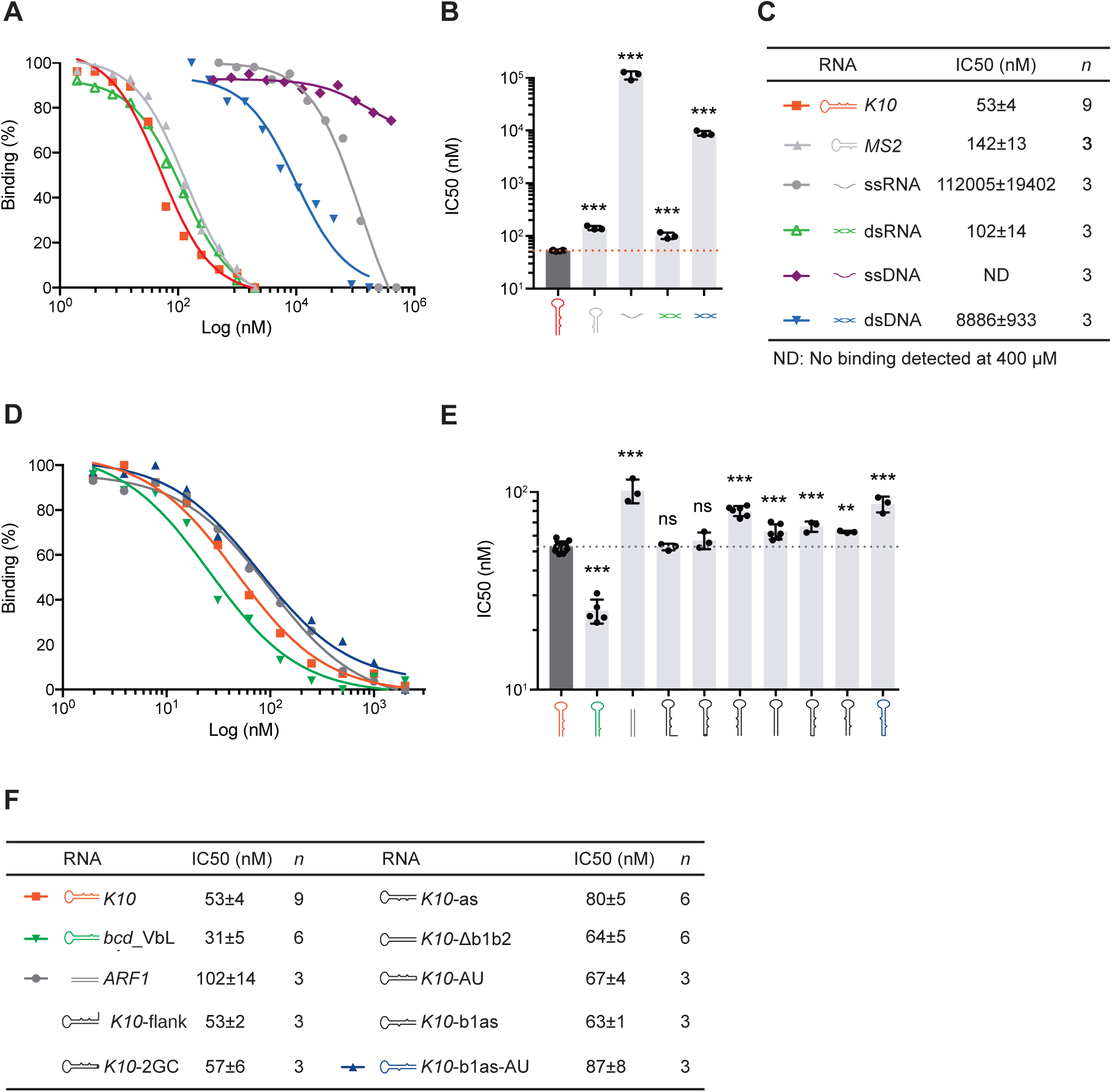
RNA binding specificity of Egl^512-819^. **(A)** and **(D)**, Egl^512-819^ was preincubated with 5′-FAM-labeled *K10* TLS RNA and unlabeled RNAs before measurement. The change in anisotropy was plotted and fitted to sigmoidal curve to obtain the IC_50_. (**B**) and (**E**), Column graph with mean IC_50_ was shown as bars, standard deviation as black error bars and individual IC_50_ values as black dots. **(C)** and **(F)**, IC_50_ values of unlabeled RNAs are shown in the table. Data are represented as mean ± SD with the number of independent measurements (n) indicated. Statistical significance was assessed using a two-sided unpaired Student’s t-test. ns, not significant; \**P* < 0.05, \*\**P* < 0.01, and \*\*\**P* < 0.001. ND, not determined.

As controls for the assay, we included another Egl target, *bcd* stem-loop 5 (*bcd*_VbL;(Lazzaretti et al., 2016)) and an Egl-independent dsRNA element from the human *ARF1* transcript(Kim et al., 2005) **(Figure 5D-F; S6D; Table S4**). *bcd*_VbL competed very efficiently with *K10* RNA, whereas *ARF1* RNA was a 1.9-fold weaker competitor.

In the structure we observed five base-directed interactions involving the G1-C48 pair, the C3-G46 pair, and A44. We focused on the G1-C48 pair because its substitution to an AU pair removes one hydrogen bond to Egl. This leads to a small but consistent (1.2-fold) reduction in competition. The structure also revealed an interaction of Egl to a bulge at C35. We inserted the bulge on the opposite strand in the same position. This *K10*-b1as RNA also showed a mild reduction in competition. When we combined the two substitutions, we found an additive effect on competition with a 1.6-fold reduction. This RNA species competes similarly to the antisense *K10* RNA, an RNA that is unable to localize in an Egl-dependent manner *in vivo*(Dienstbier et al., 2009) (**Figure 5D-F; Table S4**). Deletion of both bulges (*K10*-Δb1b2) also leads to defects in *in vivo* localization and to a 1.2-fold reduction in competition(Bullock et al., 2010).

The addition of native 10 nt flanking sequence of the RNA (*K10*-flank) and the addition of the two stabilizing base pairs, as used for structure determination (*K10*-2GC) at the base of the stem did not alter competitive ability as compared to *K10* TLS (**Figure 5D-F; Table S1**), this aligns with previous *in vivo* findings showing that such G-C substitutions do not disrupt K10 mRNA localization. In summary, sequence- and shape-specific interactions contribute to the preference of Egl^512–819^ for different RNA elements. However, the relatively efficient competition by some non-target structured RNAs indicates that the isolated RNA-binding domain is insufficient to reproduce the high selectivity observed *in vivo*. High-fidelity sorting may additionally depend on cooperative recognition of multiple RNA elements by the complete Egl–BicD complex and on other features of the cellular RNP.

### RNA binding is required for Egl function *in vivo*

We next evaluated the *in vivo* significance of RNA-protein interactions defined by our structure. We first used CRISPR/Cas9-mediated homology directed repair ((Port et al., 2014); **Figure S7A**) to create an allele of the endogenous *Drosophila egl* gene coding for a protein with the R753E, K754E and K755E mutations in the linker (*egl^L-triple^*). As described above, the wild-type residues contact in *K10* TLS and the presence of all three mutations lowers the affinity of Egl^512-819^ for the *K10 TLS in vitro* by approximately one order of magnitude (**Figure 4F-H; Table S2**).

Null mutations in *egl* have been described previously, and are most commonly associated with premature termination of translation. Flies homozygous for these alleles survive to adulthood but females are infertile. These females lack eggs due to a failure to differentiate an oocyte(Carpenter, 1994; Mach and Lehmann, 1997; Theurkauf et al., 1993). Homozygous mutant *egl^L-triple^* females were also infertile and had a complete absence of egg production (**Figure S7**). Immunostaining of ovarioles for a marker of the oocyte (Orb; (Lantz and Schedl, 1994)) and the synaptonemal complex (Corolla;(Page and Hawley, 2001) revealed the *egl^L-triple^* mutation, like previously existing *egl* null mutations, prevented differentiation of the oocyte within the germarium (**Figure 6A**). An indistinguishable phenotype was observed in ovarioles of *egl^L-triple/^egl^null^* females (**Figure 6A**). The finding that the phenotype associated with the *egl^L-triple^* allele was not complemented by an *egl^null^* chromosome generated in an independent study reveals that the phenotype is not due to an off-target mutation. Immunostaining of egg chambers with an antibody that specifically recognises Egl(Mach and Lehmann, 1997) showed that the *egl^L-triple^* mutation does not destabilise the Egl protein (**Figure S7**). Collectively, these observations reveal that the *egl^L-triple^* mutation strongly disrupts the function of the Egl protein *in vivo*.

**Figure 6.**
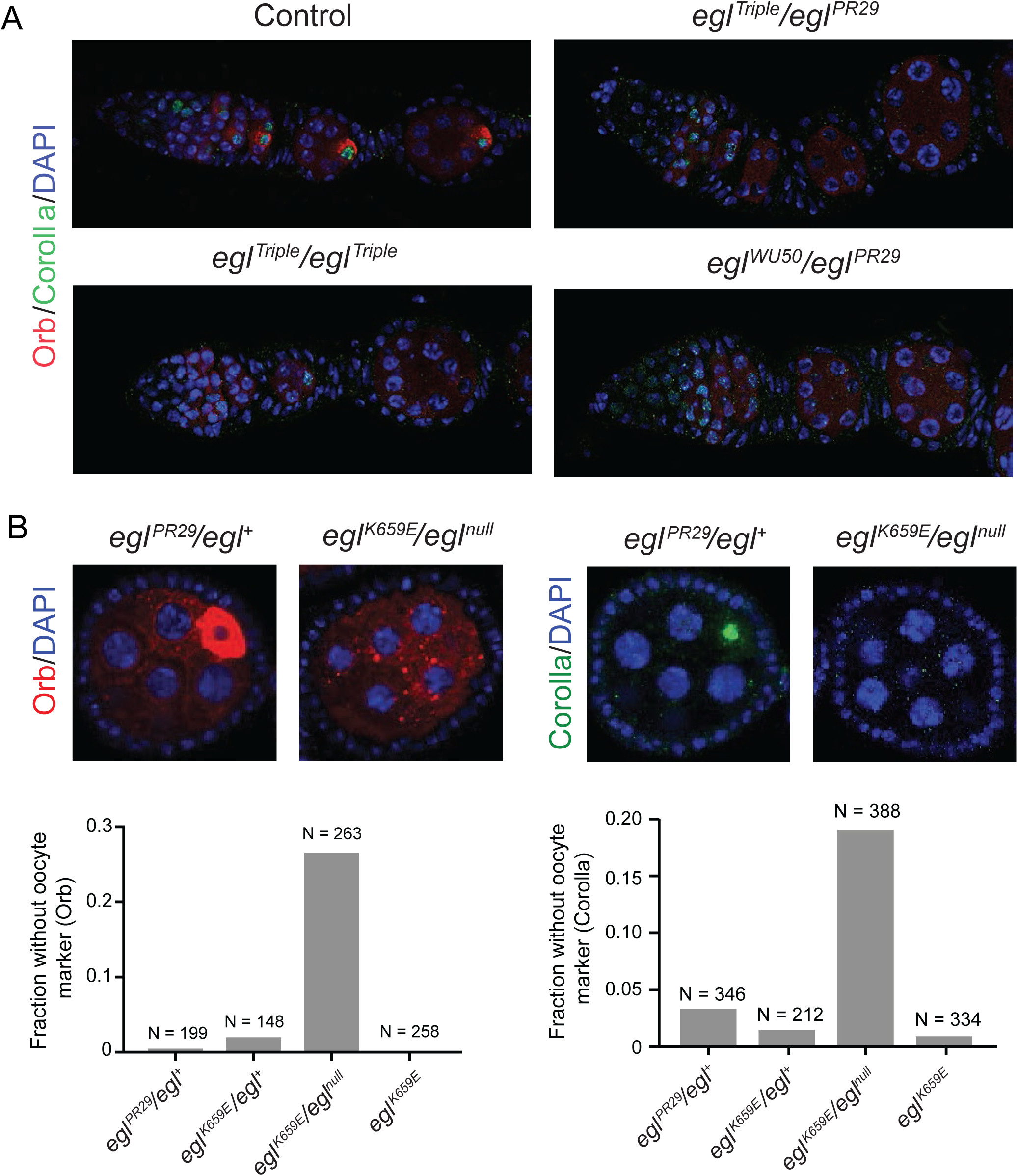
Egl residues required for RNA binding *in vitro* are required for function *in vivo*. **(A)** Images of germaria and early-stage egg chambers for the indicated genotypes stained for the oocyte markers Orb (red; cytoplasmic marker) and Corolla (green; marker of the meiotic synaptonemal complex), as well as DNA (blue; DAPI). Arrowheads label oocytes in the control. In each of the mutant genotypes, all egg chambers fail to differentiate an oocyte (images are representative of at least 200 ovarioles visualized per genotype). *PR29* and *WU50* are previously characterized *egl* null alleles(Mach and Lehmann, 1997). **(B)** Left: stage 4/5 egg chambers of the indicated genotypes stained for Orb, Corolla and DNA, illustrating the partially penetrant oocyte differentiation defect in *K659E/PR29* ovarioles. Right: quantification of oocyte differentiation phenotypes in the indicated genotypes (oocytes scored using Orb immunostaining; n = number of egg chambers examined). Control genotypes in the images in A and B are *L-Triple/+* and *PR29/+*, respectively, which had phenotypes indistinguishable from the wild-type. In B, the same control egg chamber is shown for Orb and Corolla staining. Scale bars in A and B = 20 μm.

To further examine the physiological relevance of our *in vitro* results, we used the same CRISPR methodology to introduce the K659E mutation into Egl (**Figure 6B**). As described above, K659 resides in EXO domain and contacts A44 of the *K10 TLS* in the crystal structure. K659E lowers the affinity of Egl^512-819^ for the *K10 TLS in vitro*, although to a lesser degree than the triple mutation R753E/K754E/K755E (∼3-fold vs ∼10-fold (**Figure 4**)). Females homozygous for the *K659E* mutation were fertile and exhibited no overt defects in oogenesis (**Figure 6B**). *egl^K659E/^egl^null^*females were also fertile. However, ∼30% of early egg chambers from these animals did not restrict the oocyte marker to a single cell, whereas ∼20% did not enrich the synaptonemal complex marker in a single cell (**Figure 6B**). These observations indicate a failure to specify or maintain an oocyte in a subset of cases. There was no overt decrease in Egl protein levels in the mutant egg chambers (**Figure S7**). Collectively, these data show that K659E partially disrupts the function of the Egl protein *in vivo*.

In summary, our phenotypic analysis reveals that the RNA binding residues in Egl^512-819^ are also important for the function of the full length Egl protein *in vivo.* The correlation between the extent to which the mutations we tested inhibit RNA binding *in vitro* and the degree of disruption of oocyte differentiation *in vivo* further supports the physiological relevance of our *in vitro* studies.

## Discussion

Here, we solved the X-ray crystal structure of the RNA-binding region of Egl in complex with the *K10* transport and localization signal (TLS). The structure revealed how Egl recognizes the *K10* TLS by a combination of backbone, shape-specific and base-directed interactions.. Our structure-based *in vivo* mutagenesis in *Drosophila* showed that key residues involved in *K10* TLS recognition are also important for *in vivo* function.

The EXO domain of Egl contains the conserved signature of DEDDy proteins consisting of five highly conserved residues (**Figure S1 and S4**) and shares the highest structural similarity with RNase D. RNase D is an active 3′ to 5′ DEDDy-group exonuclease with a function in the maturation of RNAs(Li et al., 1998). Our structural analysis of Egl reveals that the DEDDy residues are appropriately positioned with respect to each other in a putative catalytic centre. Remarkably, mutations of the active site residues do not affect binding to mRNA localization signals and the *in vivo* function during oogenesis(Dienstbier et al., 2009; Navarro et al., 2004). The evolutionary conservation of the DEDDy motif indicates that Egl may have exonuclease activity that contributes to other aspects of its function. However, Egl’s major function in mRNA localization appears to have evolved in parallel to the exonucleolytic activity.

The origin of new RNA-binding proteins from enzymes involved in RNA metabolism seems to have occurred repeatedly in evolution(Lazzaretti et al., 2016). For example, the RNA-binding protein Exuperantia (Exu) binds RNA by a modified exonuclease-like domain and a SAM domain. The exonuclease active-site is lost in *Drosophila* Exu but is present in some mollusc and vertebrate Exu homologs(Lazzaretti et al., 2016). Another example is Exd1, a mammalian protein related to Egl that recruits small RNAs in the piRNA pathway through its inactive exonuclease-like domain(Yang et al., 2016). In Egl, RNA binds around the catalytic site and a positively charged surface formed by the helical linker and the EHD domain. In Exu, RNA-binding has been mapped by MS cross-linking and also localizes to a patch around the active site and the SAM domain(Lazzaretti et al., 2016). In both Egl and Exu, novel RNA-binding activity may have evolved by the addition of positively charged surfaces to stabilize the interactions between the RNA substrate and the exonuclease.

Previous biochemical and single-molecule imaging studies indicate that two Egl molecules form a complex with one mRNA containing a single stem-loop structure(McClintock et al., 2018; Sladewski et al., 2018). In our SEC-RALS experiment and crystal structure, the minimal RNA-binding region of Egl binds to *K10* TLS with a 1:1 stoichiometry. This difference could be due to the shorter RNA or truncated Egl used in our study. Recent cryo-EM structures show that dimeric BicD recruits two Egl molecules and organizes two RNA-recognition sites. Each recognition site engages one structured RNA element, whereas the two sites can simultaneously recognize two elements within the same full-length transcript. Thus, the 2:1 Egl: RNA-transcript ratio reflects higher-order assembly on the dimeric BicD scaffold, while our structure defines the fundamental 1:1 interaction between the Egl RNA-binding domain and a single RNA element. Because Egl^512-819^ was studied in the absence of dimeric BicD and lacks the additional Egl regions required for assembly of the complete complex, it is not expected to reproduce the stoichiometry of the full transport RNP. Our crystallographic analysis and the cryo-EM structures are therefore complementary, defining the local RNA-recognition interface and its organization within the larger transport complex, respectively (Singh et al., 2026). The cryo-EM structures also suggest that Egl dimers couple transport initiation to coincident recognition of two structured RNA elements within one transcript.

Our crystal structure and *in vitro* affinity measurements provided insight into how Egl recognizes a localization element. Although Egl is also able to bind to ssRNA, dsRNA and other stem-loop RNA structures, the affinity of binding was higher for *K10* TLS. This suggests that Egl can discriminate RNA secondary structure and shows some sequence specificity. Several lines of evidence indicate that *in vivo* Egl binds stem-loop structures with bulges in localizing RNAs. First, the *K10* TLS is required for Egl binding and localization of the full-length mRNA (Cohen papers, Bullock 2001; 2010). Second, several mRNAs that depend on Egl for minus-end-directed transport during oogenesis and embryogenesis have similar predicted structures containing bulged regions(Bullock et al., 2010, 2003; Cohen et al., 2005; dos Santos et al., 2008; Macdonald and Kerr, 1998; Serano and Cohen, 1995; Snee et al., 2005; Van De Bor et al., 2005).

The Egl-*K10* TLS structure and our biochemical and *in vivo* experiments revealed the structural details and relevance of Egl-stem-loop interactions. We identified specific interactions between Egl and *K10*-TLS-Bulge1 that create a unique electrostatic signature for protein recognition. The importance of bulges in stem-loop elements has also been described for other RNA-protein interactions(Hermann and Patel, 2000). In the genomic RNA of HIV-1 there are bulged adenine residues that provide recognition sites for viral proteins(Ennifar et al., 1999). These interactions depend on the secondary structure or shape of the RNAs, rather than their primary sequence. Mutation of the residues in Egl that mediate interactions to Bulge1 render Egl unable to exert its function in *Drosophila* oogenesis. In addition, there are also residues of Egl that mediate base-directed interactions and backbone interactions to the stem of the RNA stem-loop. Mutations of these residues also reduced binding and *in vivo* function. These results indicate that the recognition of stem-loop structures is due to a combination of base- and shape-specific interactions.

## Accession codes

The coordinates and structure factors have been deposited in the Macromolecular Structure Database of European Bioinformatic Institute (EBI) with ID code 9UJU, 9UJY and 9UUG for Egl^512-819^-*K10* TLS complex, MBP-(GSAM)-L-EHD and *K10*-TR.

## Acknowledgements

We wish to thank the MPI-Tuebingen Crystallization Facility. We also thank the staff at the Swiss Light Source synchrotron for assistance during data collection, Claire Basquin for assistance with MALLS measurements and Stefanie Wachter for initial stages of the project. We thank G. Jékely and A. Cook for discussion and critical reading of the manuscript. This project has received funding from the European Research Council under the European Union’s Seventh Framework Programme (FP7/2007-2013), ERC grant agreement n° 310957 and the Deutsche Forschungsgemeinschaft (FOR2333 to F.B.). Li Jin is supported by MRC funding to Simon Bullock (file reference number MC_U105178790). Z.H is supported by A*STAR, National Medical Research Council (NMRC grant number MOH-001524).

## Author Contributions

Biochemical, biophysical and crystallization work was done by Z.H. and J.M.; *Drosophila* work was done by L.J.; F.B. supervised the project. All authors wrote the paper.

## Methods

### Protein expression and purification

To produce the Egl protein for crystallization and FA, Egl from *D. melanogaster* was cloned into a pET-MCN vector(Diebold et al., 2011) with an N-terminal glutathione S-transferase (GST) tag followed by a TEV protease cleavage site. Mutants and truncations were generated by site-directed mutagenesis.

GST-Egl constructs were expressed *E. coli* BL21 (*DE3*) Gold (Life Technologies) cells in TB medium at 37 °C until an OD600 of 0.6-0.8 and induced at 20 °C with 0.5 mM IPTG overnight. Cells were harvested (5000 x g, 20 min, 4 °C), resuspended in buffer A (20 mM Tris-HCl, pH 7.5, 1 M NaCl, 10% glycerol, 1 mM DTT), and lysed by homogenization. The lysate was cleared by centrifugation (18000 x g, 1 hour, 4 °C), and the supernatant was incubated with Glutathione Agarose 4B resin (Macherey-Nagel). Bound protein was washed with buffer A and buffer B (20 mM Tris-HCl, pH 7.5, 300 mM NaCl, 10% glycerol, 1 mM DTT), eluted with buffer C (20 mM Tris-HCl, pH 7.5, 300 mM NaCl, 10% glycerol, 1 mM DTT, 50 mM reduced glutathione). The protein was applied to Heparin HP Column (GE Healthcare), eluted with a gradient from 300 mM to 1 M NaCl and further purified by SEC on a HiLoad 16/600 Superdex 200 pg column (GE Healthcare) in buffer B.

The Egl^512-819^ construct was expressed and purified as described above, except that the GST-tag was removed by TEV protease overnight at 4 °C. The Egl^512-819^-*K10* TLS complex was assembled by incubating Egl^512-819^ with a 1.2-M excess of *K10* TLS on ice overnight and further purified by SEC on a HiLoad 16/600 Superdex 200 pg column (GE Healthcare) in buffer B.

The MBP-(GSAM)-L-EHD (Egl^743-819^) construct was expressed *E. coli* BL21 (*DE3*) Gold (Life Technologies) cells in autoinduction ZY medium at 37 °C until OD_600_ = 2.0, and induced at 20 °C overnight. Cells were harvested and lysed as described above. The supernatant was incubated with Amylose Resin (NEB). Bound protein was washed with buffer A and buffer B (20 mM Tris-HCl, pH 7.5, 300 mM NaCl, 10% glycerol, 1 mM DTT), eluted with buffer D (20 mM Tris-HCl, pH 7.5, 300 mM NaCl, 10% glycerol, 1 mM DTT, 50 mM Maltose) and further purified by SEC on a HiLoad 16/600 Superdex 200 pg column (GE Healthcare) in buffer E (20 mM Tris-HCl, pH 7.5, 150 mM NaCl, 1 mM DTT, 10 mM Maltose).

### Crystallization, data collection and analysis

Crystallization conditions were initially screened at 4°C and at 20°C using a Mosquito nanodrop dispenser (TTP Labtech) in 96-well sitting-drop plates with 100 nl protein plus 100 nl reservoir. Crystals of Egl^512-819^-*K10* TLS complex grew in 50 mM sodium citrate, pH 4.8, 30% (w/v) MPD, 10 mM CaCl_2_, 2.5 mM Spermine at 20 °C. For X-ray data collection, a single crystal was harvested without cryoprotectant and flash cooled in liquid nitrogen. Diffraction data were collected at Beamline PXII of the Swiss Light Source. The diffraction data were indexed, integrated, and scaled using the XDS program package(Kabsch, 2010). A weak molecular replacement solution [TFZ=8.4] was obtained by using a trimmed poly-Ala model (RNAse D, PDB ID: 4NLB) generated by I-TASSER(Yang and Zhang, 2015). The model building of Egl^512-819^ was performed with MR-Rosetta(Terwilliger et al., 2012) and manual building in COOT(Emsley et al., 2010), and the model of *K10* TLS was built using Brickworx(Chojnowski et al., 2015) and manual run in COOT(Emsley et al., 2010). Refinement was carried out in Refmac5 using TLS and jelly body(Vagin et al., 2004).

Crystals of MBP-(GSAM)-L-EHD grew in 100 mM Tris-HCl, pH 7.5, 18% (w/v) PEG 6000, 50 mM MgCl_2_ at 4 °C. For X-ray data collection, a single crystal was transferred to a cryoprotectant solution with mother liquor supplemented with addition of 20% glycerol, and flash cooled in liquid nitrogen. Diffraction data were collected and processed as described above. The structure was solved by molecular replacement with PHASER(McCoy et al., 2007) using the structure of maltose binding protein (PDB ID: 1ANF)(Quiocho et al., 1997) as the search model. All subsequent model building was performed in COOT(Emsley et al., 2010) and restrained refinement in Phenix(Terwilliger et al., 2012).

Protein structure figures were generated using PyMOL (Schrödinger, LLC.). The quality of the final models was validated with the wwPDB Validation Server (https://validate-rcsb.wwpdb.org/). Interaction surfaces were calculated with PISA(Krissinel and Henrick, 2007).

### Fluorescence anisotropy

For direct binding assay, measurements were carried out using an Infinite F200 plate reader (Tecan) at 20°C. 5’-6-FAM-labeled RNA was added to each reaction at a fixed concentration of 10 nM with different concentrations of purified Egl truncations or mutants in 50 µl buffer F (20 mM HEPES-NaOH pH 7.5, 100 mM NaCl, 10% Glycerol). Each titration point was measured three times, with an integration time of 40 µs, with 485 nm and 535 nm as the excitation and emission wavelength, respectively. The data were analyzed using the Hill equation with the Prism 6 software (GraphPad).

For competition binding assay, 150 nM Egl^512-819^ was preincubated with 10 nM 5’-FAM-labeled *K10* TLS RNA for 10 min at 20°C and further incubated with different concentrations of unlabeled RNAs for 10 min at 20°C in 50 µl buffer F before measurement. The change of anisotropy was plotted and fitted to sigmoidal curve to obtain the IC_50_ with Prism 6 (GraphPad).

The sequences of the RNA and DNA oligonucleotides (Integrated DNA Technologies) are shown as follows (5’ to 3’):

*K10* (CUUGAUUGUAUUUUUAAAUUAAUUCUUAAAAACUACAAAUUAAG);

*K10*-Δb1b2 (CUUGAUUGUAUUUUUAAAUUAAUUCUUAAAAAUACAAUUAAG);

*K10*-as (GAAUUAAACAUCAAAAAUUCUUAAUUAAAUUUUUAUGUUAGUUC);

*K10*-b1as (CUUGAUUGUACUUUUUAAAUUAAUUCUUAAAAAUACAAAUUAAG);

*K10*-AU (AUUGAUUGUAUUUUUAAAUUAAUUCUUAAAAACUACAAAUUAAU);

*K10*-b1as-AU (AUUGAUUGUACUUUUUAAAUUAAUUCUUAAAAAUACAAAUUAAU);

*K10*-2GC (GGCUUGAUUGUAUUUUUAAAUUAAUUCUUAAAAACUACAAAUUAAGCC);

*K10*-flank (CUUGAUUGUAUUUUUAAAUUAAUUCUUAAAAACUACAAAUUAAGAUCACUCUGU);

*ARF1* sense (dsRNA sense) (UGAGUGCCAGAAGCUGCCUC);

*ARF1* antisense (GAGGCAGUUUCUGGUACUCA);

dsRNA antisense (GAGGCAGCUUCUGGCACUCA);

ssRNA (polyU20) (UUUUUUUUUUUUUUUUUUUU);

ssRNA (polyAU20) (AUAUAUAUAUAUAUAUAUAU);

*bcd*_VbL (long) (GGGCCCAAAAUGAAAAAUGUUUCUCUUGGGCGUAAUCUCAUACAAUGAUUACCCUUAAAGAUCGAACAUUUAAACAAUAAUAUUUGGG);

*bcd*_VbS (short) (AAAUGUUUCUCUUGGGCGUAAUCUCAUACAAUGAUUACCCUUAAAGAUCGAACA UUU);

ssDNA (dsDNA sense) (AGGCAGTTTCTGGTACTCAG);

dsDNA antisense (CTGAGTACCAGAAACTGCCT);

*MS2* (ACAUGAGGAUUACCCAUGU).

### Right angle light scattering (RALS)

Samples were loaded onto a Superdex 75 5/150 column (GE Healthcare) connected to a TDA302 detector array (Viscotek), in buffer B at room temperature. Chromatography runs were performed with 2 mg/ml of purified Egl alone, or pre-incubated with a 1.1-M excess of *K10* TLS on ice for 10 min. The sample volume was 20 µl. Data were analyzed with OmniSEC 4.5 software.

### Analytical size-exclusion chromatography

Egl^512-819^ (32 uM, 500 ul) alone, or pre-incubated with a 1.2-M excess of *K10* TLS were loaded on a Superdex 75 10/300 GL column (GE Healthcare) in buffer B, and UV absorbance was monitored at 280 and 260 nm.

### Circular dichroism

CD spectroscopy was performed on a JASCO-810 CD system. Wavelength scans were averages of five scans between 200 nm and 250 nm collected at 20°C in a quartz cuvette with a 1-mm path length with 0.2 mg/ml protein in buffer B.

### Generation of *Drosophila egl* alleles by CRISPR/Cas9-mediated homology-directed repair

We used our previously reported CRISPR/Cas9 methodology(Port et al., 2014) to produce *egl* alleles encoding proteins with mutations of specific RNA-binding residues. Briefly, single-stranded oligonucleotide donors coding for the desired mutations (Ultramers, IDT; 500 ng/μl) were injected into the posterior of embryos that had a *nos-cas9* transgene(Port et al., 2014) (CFD2) and a transgene (integrated at attP40) that expresses a single guide (sg)RNA targeting the *egl* locus adjacent to the region to be repaired by the respective donor (embryos were generated by crossing *nos-cas9* females with sgRNA males). Spacer sequences for the two sgRNAs (*sgRNA^L-Triple^* and *sgRNA^K659E^*) had previously been cloned via complementary oligonucleotides into the transformation vector pCFD3(Port et al., 2014), which expresses sgRNA from the *U6:3* promotor. CRISPR RGEN tools (http://www.rgenome.net/) revealed the selected sgRNAs have no other predicted targets in the *Drosophila* genome. Following injection of the donor into the transgenic embryos, flies heterozygous for the desired mutation were identified by Sanger sequencing of PCR products, as described previously(Port and Bullock, 2016). Sequences of spacer and donor oligos, as well as oligos used for sequencing the targeted region, are provided in Table S1. Balanced mutant stocks were established from the offspring of positive animals, followed by generation of homozygous females for phenotypic analysis.

### Immunostaining of ovarioles

Ovarioles were dissected in PBS with 0.5% Tween (PBT0.5) and fixed in 4% paraformaldehyde/PBT0.5 for 20 minutes. Following washing in PBT0.5 (3 x 5 min and 4 x 15 min), samples were blocked in 20% Western Blocking Reagent (Roche)/PBT0.5 (Blocking Buffer) for 1 h. The following primary antibodies were then used overnight at 4°C in Blocking Buffer: (1) a combination of mouse anti-Orb (mixture of clones 4H8 and 6H4(Lantz et al., 1994) (Developmental Studies Hybridoma Bank; each diluted 1:200)) and rabbit anti-Corolla(Collins et al., 2014) (gift of Scott Hawley, Stowers Institute (diluted 1:2000)), or (2) rabbit anti-Egl (Mach and Lehmann, 1997)(gift of Ruth Lehmann, Skirball Institute; diluted 1:3000)). Following 3 x 5 min and 4 x 15 min PBT0.5 washes, secondary antibodies in Blocking Buffer (Alexa488-conjugated donkey-anti-rabbit (Life Technologies (A21206)); Alexa555-conjugated donkey-anti-mouse (Life Technologies (A31570) (each diluted 1:500)) were added for 2 h at room temperature. Washing steps were repeated and, after 2 brief washes in PBS, ovarioles were mounted in Vectashield containing DAPI (VectorLabs). Ovarioles were imaged with a Zeiss 780 confocal microscope using a Zeiss 40 x 1.3NA oil objective (except for Figure S7A (Zeiss 20 x 0.8NA objective)).

---

**Figure S1 related to Figure 1.**
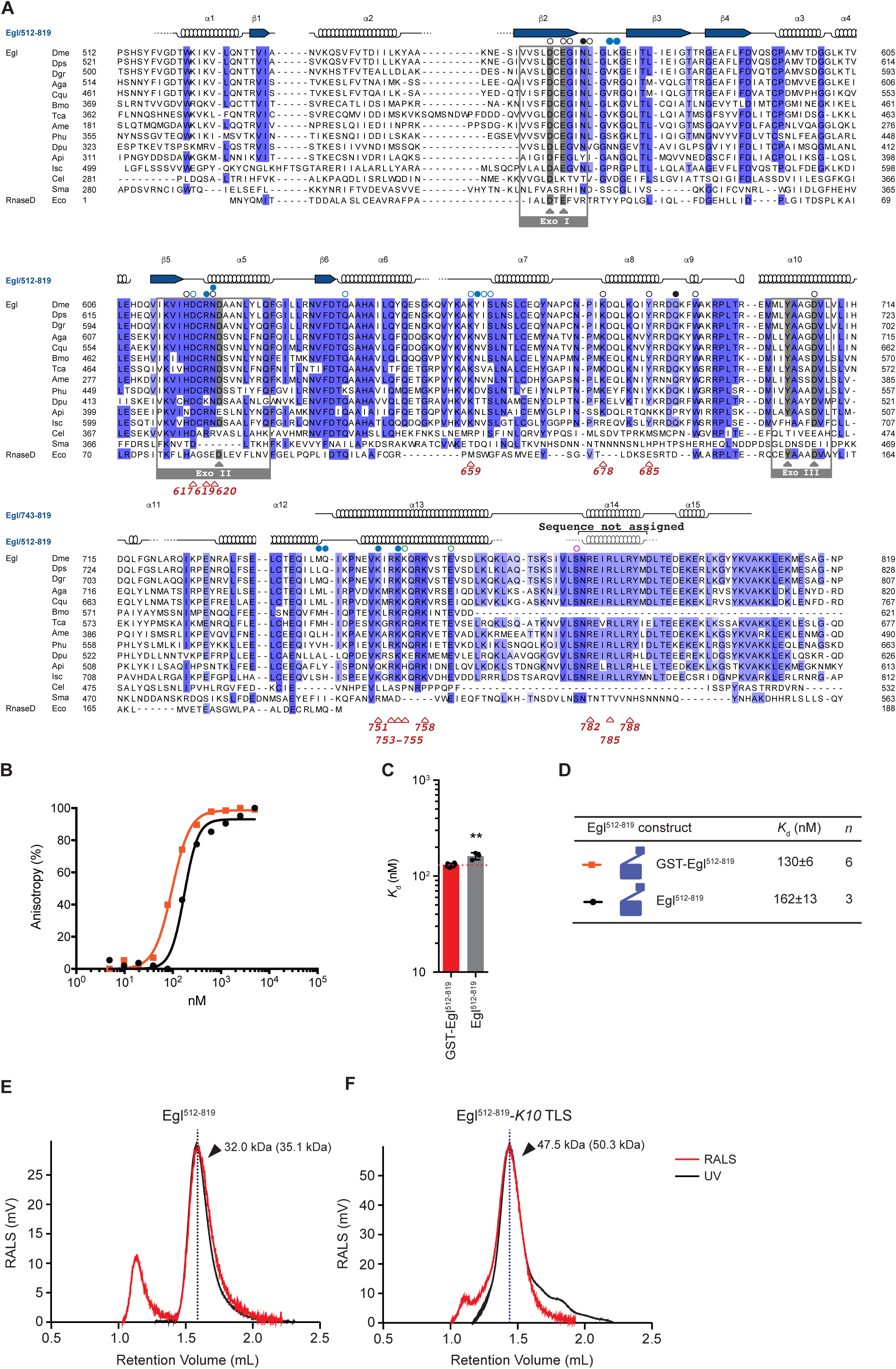
Alignment of Egl homologs and Egl RNA binding region. **(A)** Sequences used to generate the alignment include: [Insecta, order Diptera] *D. melanogaster* (Dme), *D. sechellia*, *D. simulans*, *D. erecta*, *D. yakuba*, *D. ananassae*, *D. persimilis*, *D. pseudoobscura pseudoobscura*, *D. willistoni*, *D. virilis*, *D. mojavensis*, *D. grimshawi*, *A. gambiae* (Aga), *C. quinquefasciatus*; [Lepidoptera] *B. mori* (Bmo), *H. melpomene*, *D. plexippus*; [Coleoptera] *T. castaneum* (Tca); [Hymenoptera] *A. cephalotes*, *A. mellifera* (Ame); [Phthiraptera] *P. humanus corporis*; [Hemiptera] *A. pisum* (Api); [Crustacea] *D. pulex* (Dpu); [Chelicerata] *I. scapularis* (Ixo); [Nematoda] *C. elegans* (Cgi), *C. briggsae*, *P. pacificus*; [Plathyelmintes] *S. mansoni* (Dre); [Echinodermata] *S. purpuratus*. Numbering refers to the Dme sequence. The secondary structure of Dme Egl^512-819^ and L-EHD is schematized above the alignment (beta-sheets: 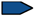; alpha-helices: 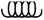; residues not ordered in the structure are shown as dotted lines; gaps indicate regions for which no structural information is available). Residues interacting with the 5′ or 3′ strand of the *K10* stem-loop RNA are marked with a filled or with an empty dot above the Dme sequence, respectively; conserved residues are highlighted in blue according to 80% sequence identity in Jalview(Waterhouse et al., 2009). Catalytic residues in the exonuclease domain: 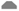; mutated residues: 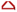; conserved catalytic residues are highlighted in gray. The alignment was generated with MUSCLE(Edgar, 2004), visualized with Jalview (Waterhouse et al., 2009) and edited in Adobe Illustrator. **(B)** RNA binding properties of GST-tagged and untagged Egl^512-819^. *K*_d_ values were determined by FA. 5′-FAM-labeled *K10* TLS was incubated with increasing concentrations of GST-Egl^512-819^ or Egl^512-819^ and the data were fitted to the Hill equation to obtain the *K*_d_. Data are represented as mean ± SD. **(C)** Column graph with mean *K*_d_ was shown as bars in the same color-coding, standard deviation as black error bars and individual *K*_d_ values as black dots. **(D)** *K*_d_ values of GST-Egl^512-819^ or Egl^512-819^ for *K10* TLS are shown in the table. Data are represented as mean ± SD with the number of independent measurements (n) indicated. Statistical significance was assessed using a two-sided unpaired Student’s t-test. ns, not significant; \**P* < 0.05, \*\**P* < 0.01, and \*\*\**P* < 0.001. ND, not determined. RALS profiles of Egl^512-819^ unbound **(E)** and in complex with *K10* TLS RNA **(F)**. For each plot, the calculated molecular weight (MW) at the peak is indicated in black; the predicted molecular weight is shown in parenthesis. RALS and UV curves are colored in red and black, respectively.

**Figure S2 related to Figure 2.**
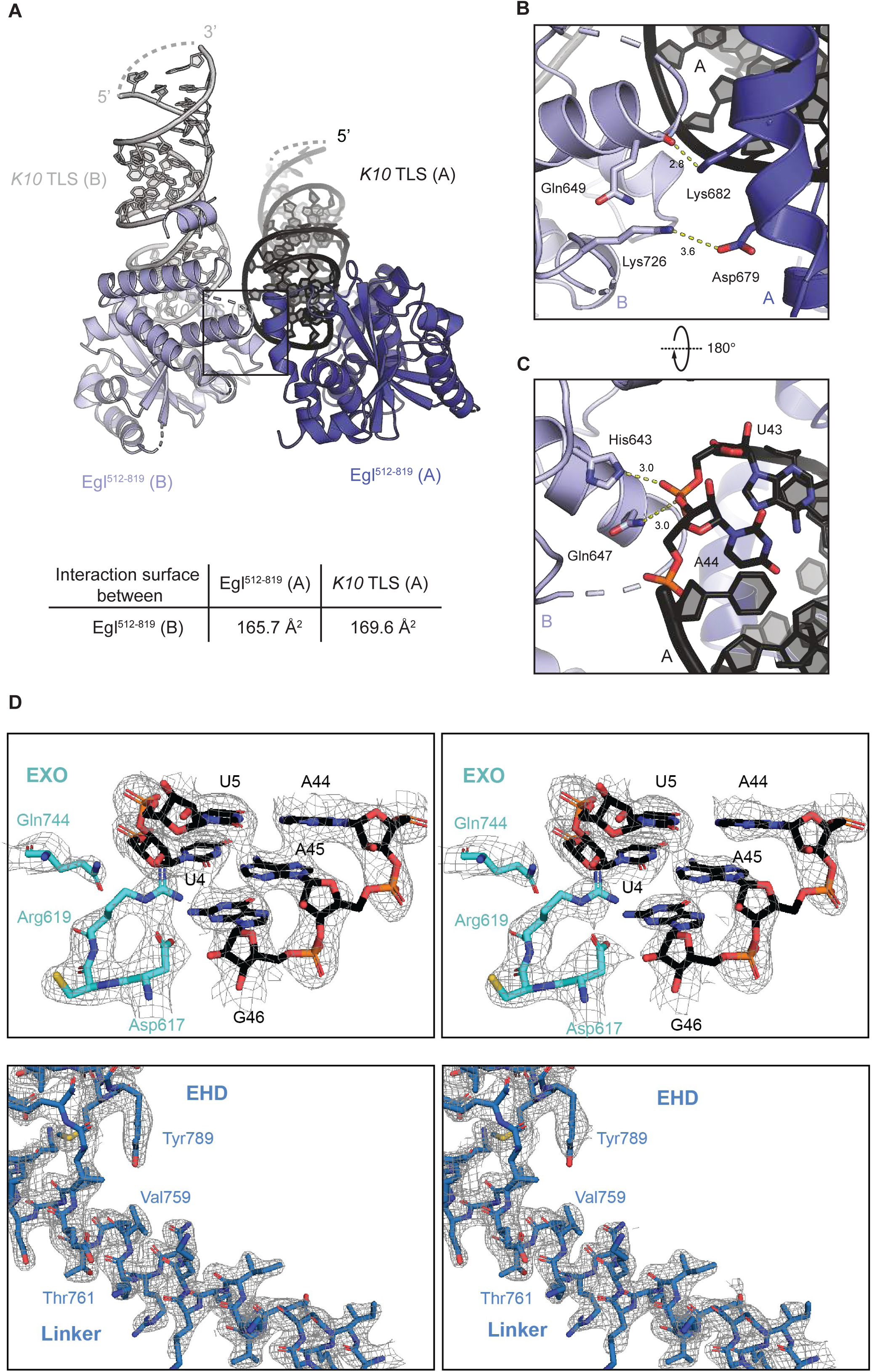
Egl^512-819^ multimerization state in the complex and quality of the electron density. **(A)** The crystal lattice of Egl^512-819^-*K10* TLS includes two complexes in the ASU. In molecule A, Egl^512-819^ is rendered as cartoon in blue, while *K10* RNA is in black. In molecule B, the protein is in light blue and the RNA in gray. A table displaying the extension of the interface between the two complexes in the ASU is shown below. **(B)** A zoomed in view of the interactions at the interface between Egl^512-819^ molecule A and Egl^512-819^ molecule B. The side chain of Lys682^A^ interacts with the backbone carboxyl of Gln649 ^B^. **(C)** Interactions between Egl^512-819^ molecule B and *K10* TLS molecule A. *K10* TLS RNA^A^ backbone contacts His643^B^ and Gln647^B^. With the exception of Lys726, all other interacting residues are evolutionarily conserved. **(D)** Stereo views of electron-density maps for the Egl^512-819^-K10 TLS complex superimposed on the final refined model. The refined 2∣Fo∣−∣Fc∣ map is shown on the left, and the composite simulated-annealing omit 2∣Fo∣−∣Fc∣ map is shown on the right in the same orientation. Both maps are contoured at 1σ.

**Figure S3 related to Figure 3.**
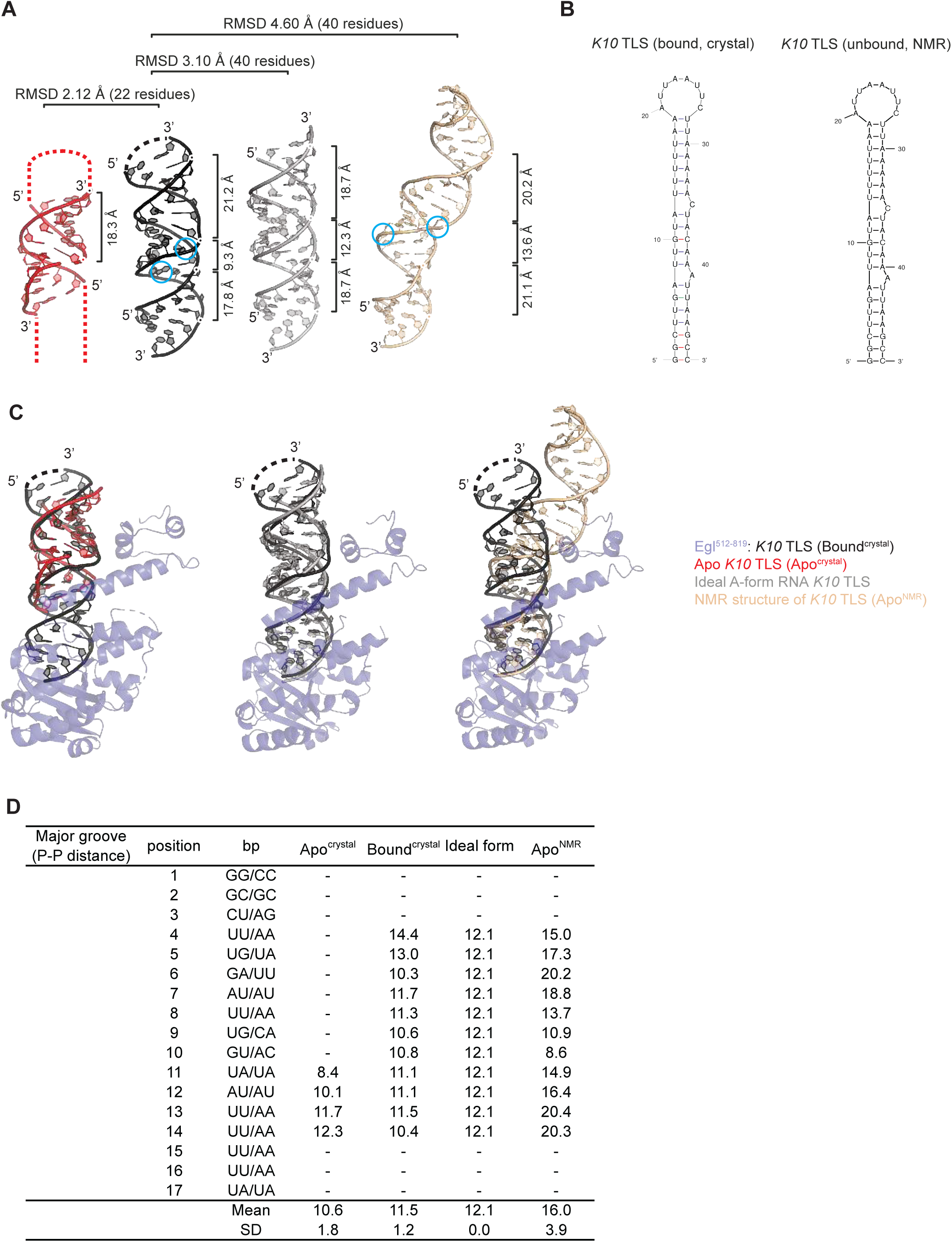
Conformational changes of *K10* TLS in the complex and in isolation. **(A)** From left to right: crystal structures of *K10* TLS (in red), of the Egl^512-819^ bound *K10* TLS (in black), ideal A-form RNA *K10* TLS lacking the two bulges C35 and A41 (Δbulges) (in gray) and NMR solution structure of *K10* TLS (PDB ID.: 2KE6; Bullock et al. 2010; in gold) in a similar view. The width of major/minor grooves and RMSD are indicated. **(B)** Schematic representation of both Egl^512-819^ bound *K10* TLS and NMR solution structure of *K10* TLS. **(C)** Structural alignments of the Egl^512-819^ bound *K10* TLS with *K10* TLS (left), with an ideal A-form RNA *K10* TLS (Δbulges) (middle) and with the NMR solution structure of *K10* TLS. **(D)** *K10* TLS structural parameters calculated with w3DNA (Zheng et al., 2009). Major groove widths: direct P-P distances, not accounting for Van der Waals radii of the phosphate groups. Graphical illustrations of the parameters are reproduced from http://x3dna.org (Lu and Olson, 2003).

**Figure S4 related to Figure 2 and 3.**
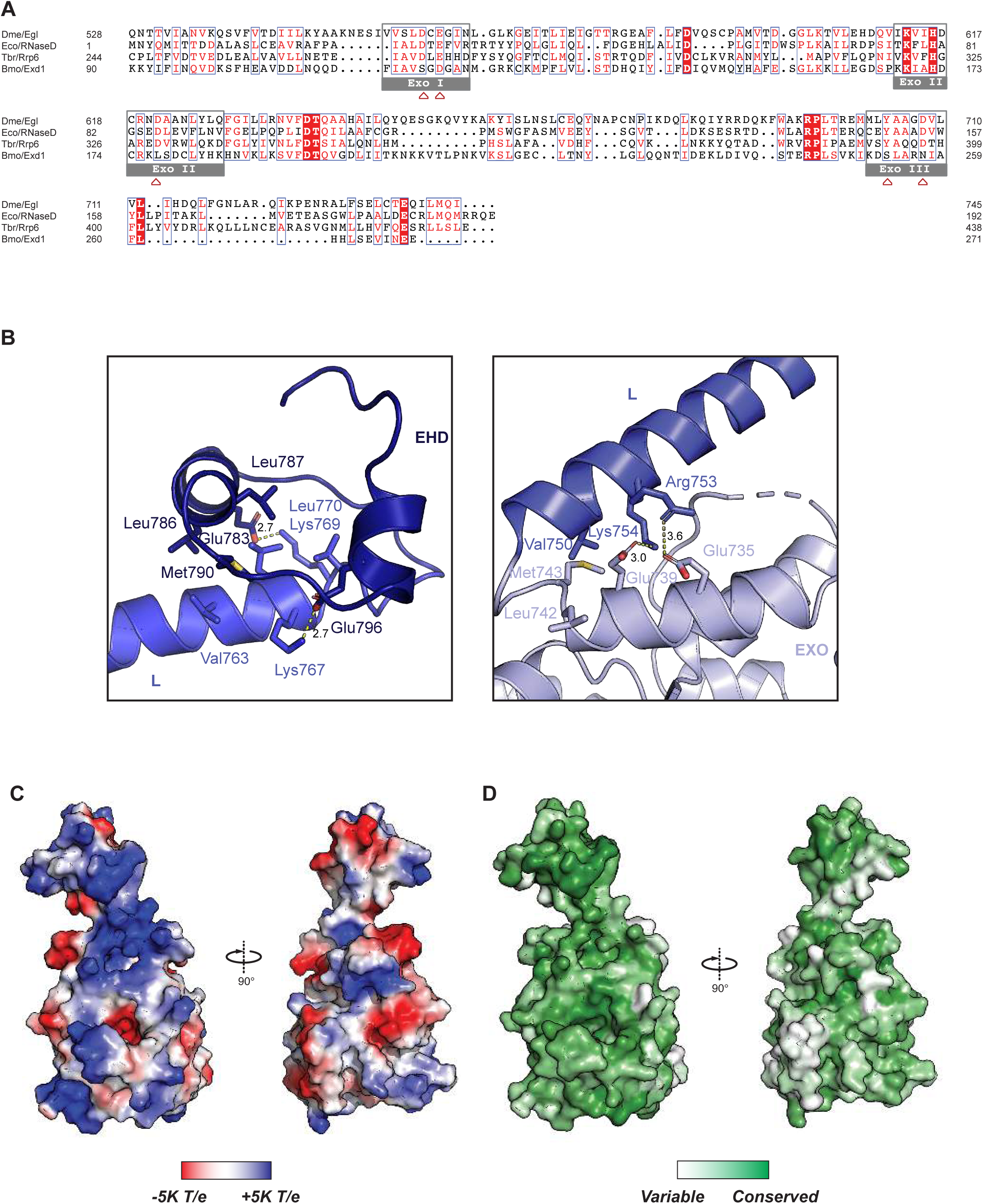
Alignment of Egl-related EXO domains, interdomain interaction of Egl^512-819^ and *K10* RNA binds at positively charged and conserved residues on the surface of Egl^512-819^. (A) Sequence alignment of the EXO domains of *D. melanogaster* Egl (Egl_Dme), *E. coli* RNase D (RNaseD_Eco), *T. brucei* RRP6 (RRP6_Tbr) and *B. mori* EXO domain-containing 1 (Exd1_Bmo). Conserved residues are highlighted in red. Green boxes indicate exonuclease signature motifs, while signature catalytic residues are marked with 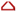. The alignment was generated with MUSCLE (Edgar, 2004), visualized with ESPript (Robert and Gouet, 2014) and edited in Adobe Illustrator. The catalytic sites are marked by light gray boxes. **(B)** Interdomain interaction of Egl^512-819^. (**C**-**D**) Surface representation of Egl in the same orientation as in **(C)** and rotated 90° along the *x* axis. In **(C)**, the surface is colored according to electrostatic potential, with positively charged residues in blue. In **(D)**, the surface is colored according to conservation, with a gradient from white to green indicating increasingly conserved residues. Conservation scores were calculated from the Egl sequences included in the multiple sequence alignment shown in **Figure S1A**, by using ConSurf (Ashkenazy 2016).

**Figure S5 related to Figure 4.**
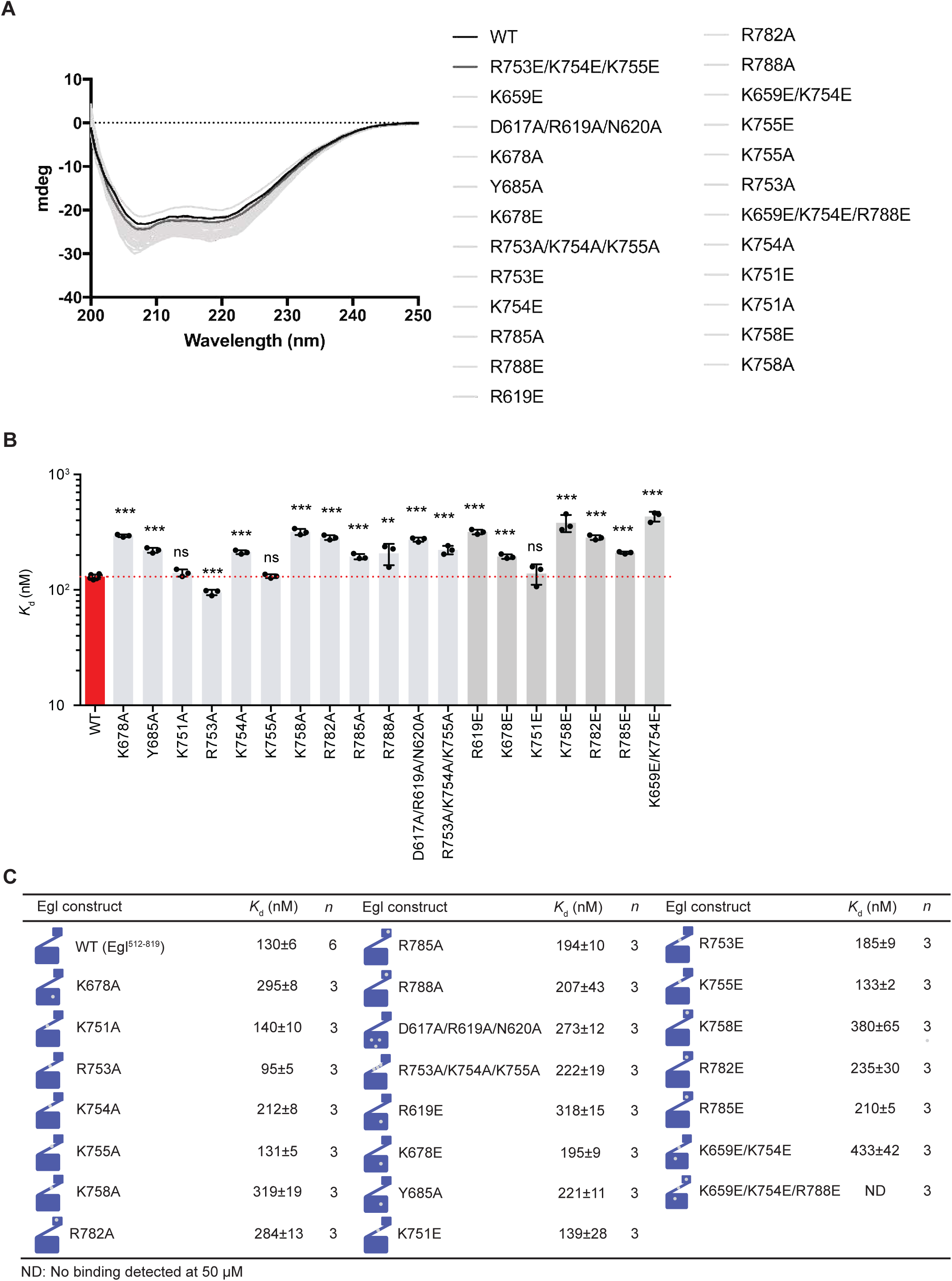
Binding affinity of Egl alanine mutants to *K10* TLS and Circular Dichroism spectra of Egl mutants used in this study. **(A)** Circular Dichroism (CD) spectra of recombinantly purified Egl mutants used in FA. Wavelength scans are averages of five scans between 200 nm and 250 nm collected at 20°C in a quartz cuvette with a 1-mm path length with 0.2 mg/ml protein in buffer (20 mM Tris-HCl, pH 7.5, 300 mM NaCl, 10% glycerol, 1 mM DTT). Wavelength scans are shown as solid curves with Egl WT colored in black, the R753E/K754E/K755E triple mutant in dark gray, and the other mutants in light gray. **(B)** Column graph with mean *K*_d_ shown as bars, standard deviation as black error bars and individual Kd values as black dots. (C) *K*_d_ values between Egl alanine mutants and *K10* TLS are shown in the table. Data are represented as mean ± SD with the number of independent measurements (n) indicated. Statistical significance was assessed using a two-sided unpaired Student’s t-test. ns, not significant; \**P* < 0.05, \*\**P* < 0.01, and \*\*\**P* < 0.001. ND, not determined.

**Figure S6 related to Figure 5.**
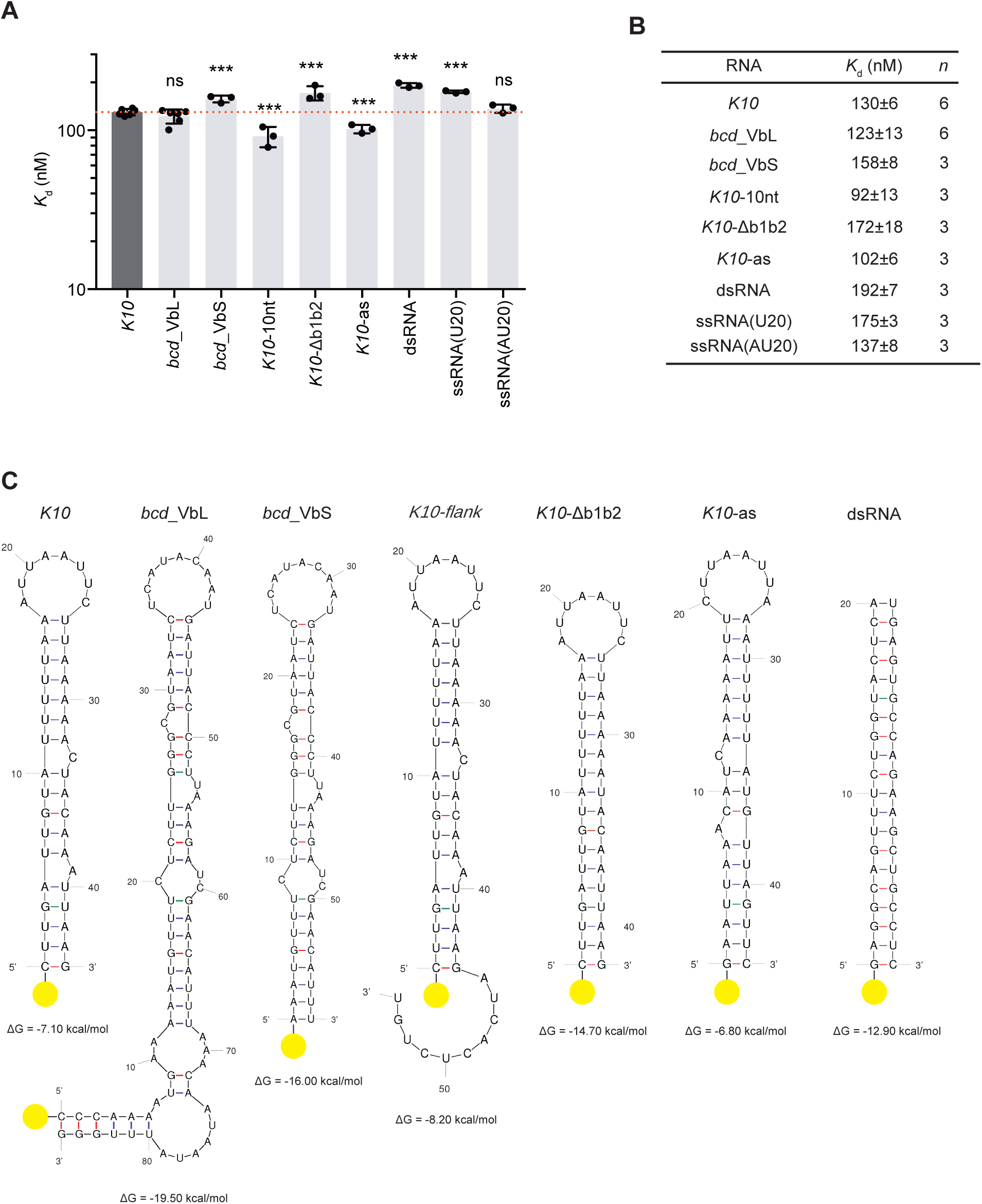
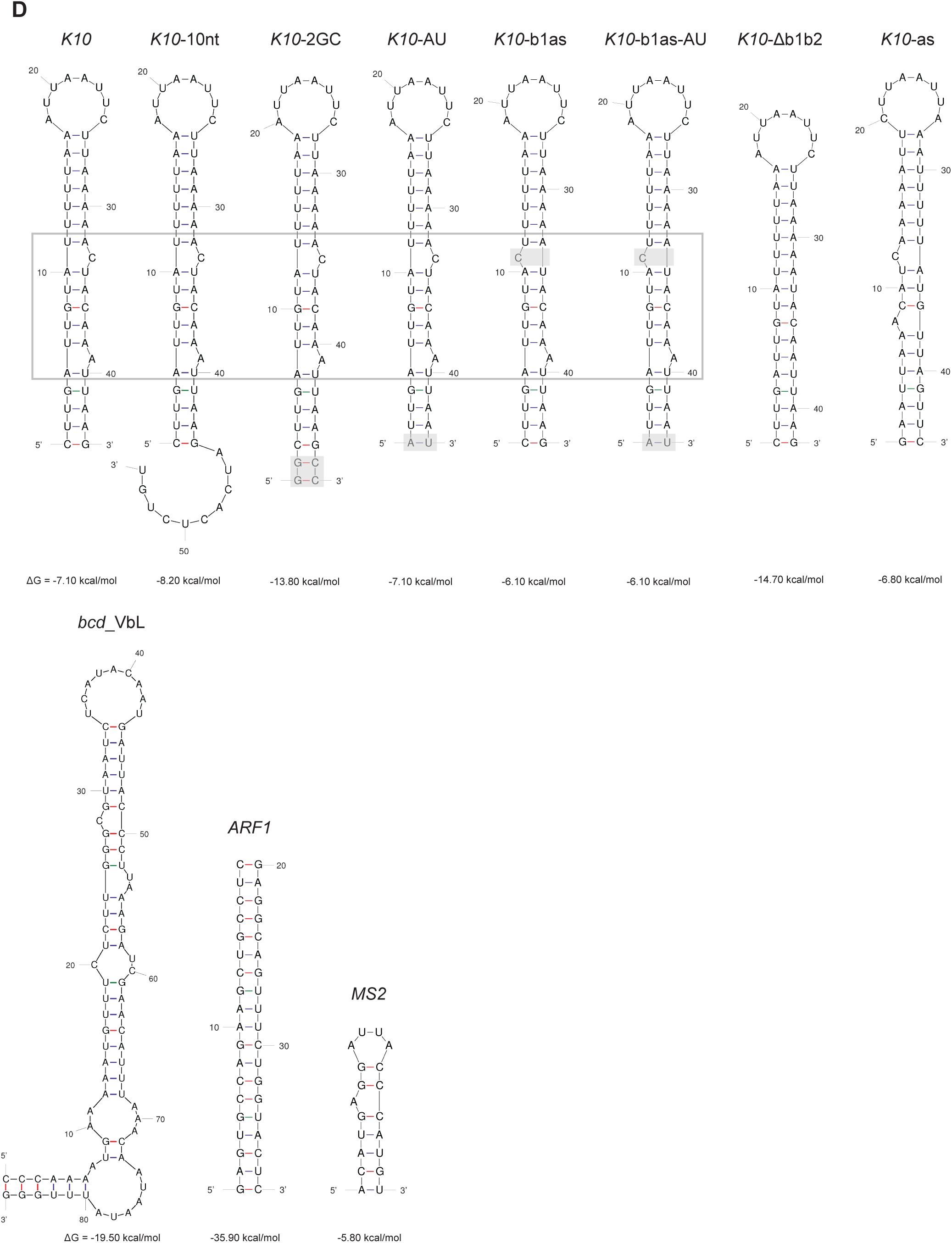
Binding affinity of Egl^512-819^ to RNA variants. **(A)** and **(B)** *K*_d_ values were determined by FA. 5′-FAM-labeled RNA variants were incubated with increasing concentrations of Egl^512-819^ and the data were fitted to the Hill equation to obtain the *K*_d_. Data are represented as mean ± SD. **(A)** Column graph with mean *K*_d_ was shown as bars, standard deviation as black error bars and individual *K*_d_ values as black dots. **(B)** *K*_d_ values between Egl^512-819^ and RNA variants are shown in the table. Data are represented as mean ± SD with the number of independent measurements (n) indicated. Statistical significance was assessed using a two-sided unpaired Student’s t-test. ns, not significant; \**P* < 0.05, \*\**P* < 0.01, and \*\*\**P* < 0.001. ND, not determined. **(C)** Secondary structures of RNA variants used for direct binding assay. **(D)** Secondary structures of RNA variants used in competition assay. Secondary structures of RNA variants were generated with mFold (Zuker 2003).

**Figure S7 related to Figure 6.**
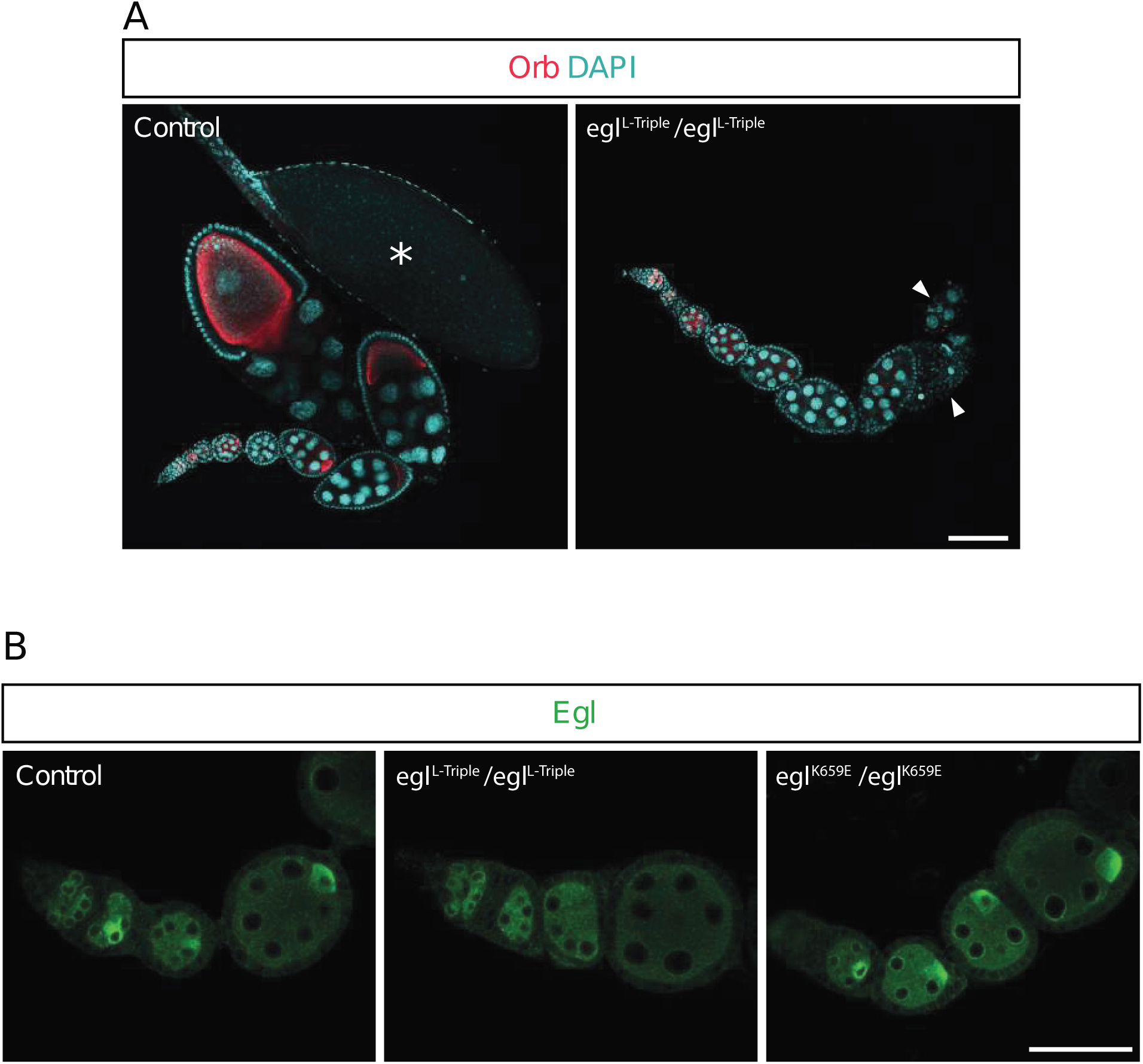
Additional characterisation of *egl* mutations in *Drosophila*. **(A)** Female flies homozygous for the *L-Triple* mutation fail to produce mature oocytes (labelled in the control by an asterisk). In the control ovariole, developing oocytes within syncytial egg chambers are visualised by staining with an anti-Orb antibody (red). DNA is labelled with DAPI (cyan). In the absence of oocyte differentiation, mutant egg chambers eventually degenerate (arrowheads), which is consistent with previous observations in *egl* null mutants(Mach and Lehmann, 1997). Scale bar, 100 μm. Images are representative of > 200 ovarioles examined per genotype. Control genotype is *PR29/*+, which has indistinguishable oocyte development from the wild-type. **(B)** The mutations in *egl* that impair RNA binding do not prevent protein expression. Images are of germaria and early egg chambers. Note that Egl (green) accumulates in the oocyte in control egg chambers by microtubule-based transport(Mach and Lehmann, 1997). This accumulation is also seen in *K659E/K659E* egg chambers, which have no overt defects in oocyte differentiation (**Fig. 6B**). In contrast, homozygous *L-Triple* fail to differentiate an oocyte (**Fig. 6A**) and therefore do not accumulate Egl asymmetrically in the egg chamber. Thus, for this genotype, protein levels in the rest of the egg chamber should be compared to control. Control is wild type (*w^1118^*). Scale bar = 50 μm. Images are representative of > 75 ovarioles imaged per genotype.

**Supplementary Table S1.** Egl512-819 versus Egl constructs of different lengths.

| Construct | Kd (nM) | n | Fold | P value | Significance |
| --- | --- | --- | --- | --- | --- |
| Egl512-819 | 130 ± 6 | 6 | 1.00 | Reference | Reference |
| Egl1-819 | 127 ± 7 | 3 | 0.98 | 0.522 | ns |
| Egl512-1004 | 144 ± 6 | 3 | 1.11 | 0.013 | * |
| Egl512-931 | 164 ± 7 | 3 | 1.26 | 1.23e-04 | *** |
| Egl512-751 | ND | 3 | NA | NA | NA |
| Egl512-770 | 807 ± 104 | 3 | 6.21 | 5.62e-07 | *** |
| Egl743-819 | 262 ± 9 | 3 | 2.02 | 2.64e-08 | *** |

**Supplementary Table S2.** Egl512-819 versus Egl mutants.

| Construct | Kd (nM) | n | Fold | P value | Significance |
| --- | --- | --- | --- | --- | --- |
| WT Egl512-819 | 130 ± 6 | 6 | 1.00 | Reference | Reference |
| K678A | 295 ± 8 | 3 | 2.27 | 3.89e-09 | *** |
| K751A | 140 ± 10 | 3 | 1.08 | 0.096 | ns |
| R753A | 95 ± 5 | 3 | 0.73 | 5.58e-05 | *** |
| K754A | 212 ± 8 | 3 | 1.63 | 4.93e-07 | *** |
| K755A | 131 ± 5 | 3 | 1.01 | 0.812 | ns |
| K758A | 319 ± 19 | 3 | 2.45 | 6.33e-08 | *** |
| R782A | 284 ± 13 | 3 | 2.18 | 3.83e-08 | *** |
| R785A | 194 ± 10 | 3 | 1.49 | 5.43e-06 | *** |
| R788A | 207 ± 43 | 3 | 1.59 | 0.002 | ** |
| D617A/R619A/N620A | 273 ± 12 | 3 | 2.10 | 4.50e-08 | *** |
| R753A/K754A/K755A | 222 ± 19 | 3 | 1.71 | 8.65e-06 | *** |
| R619E | 318 ± 15 | 3 | 2.45 | 1.89e-08 | *** |
| K678E | 195 ± 9 | 3 | 1.50 | 3.43e-06 | *** |
| Y685A | 221 ± 11 | 3 | 1.70 | 7.11e-07 | *** |
| K751E | 139 ± 28 | 3 | 1.07 | 0.447 | ns |
| R753E | 185 ± 9 | 3 | 1.42 | 1.05e-05 | *** |
| K755E | 133 ± 2 | 3 | 1.02 | 0.440 | ns |
| K758E | 380 ± 65 | 3 | 2.92 | 2.04e-05 | *** |
| R782E | 235 ± 30 | 3 | 1.81 | 4.83e-05 | *** |
| R785E | 210 ± 5 | 3 | 1.62 | 2.14e-07 | *** |
| K659E/K754E | 433 ± 42 | 3 | 3.33 | 3.20e-07 | *** |
| K659E/K754E/R788E | ND | 3 | NA | NA | NA |

**Supplementary Table S3.** K10 versus other RNAs: Kd.

| RNA | Kd (nM) | n | Fold | P value | Significance |
| --- | --- | --- | --- | --- | --- |
| K10 | 130 ± 6 | 6 | 1.00 | Reference | Reference |
| bcd_VbL | 123 ± 13 | 6 | 0.95 | 0.259 | ns |
| bcd_VbS | 158 ± 8 | 3 | 1.22 | 5.59e-04 | *** |
| K10-10nt | 92 ± 13 | 3 | 0.71 | 4.25e-04 | *** |
| K10-Delta b1b2 | 172 ± 18 | 3 | 1.32 | 9.45e-04 | *** |
| K10-as | 102 ± 6 | 3 | 0.78 | 3.04e-04 | *** |
| dsRNA | 192 ± 7 | 3 | 1.48 | 2.34e-06 | *** |
| ssRNA(U20) | 175 ± 3 | 3 | 1.35 | 6.48e-06 | *** |
| ssRNA(AU20) | 137 ± 8 | 3 | 1.05 | 0.179 | ns |

**Supplementary Table S4.**
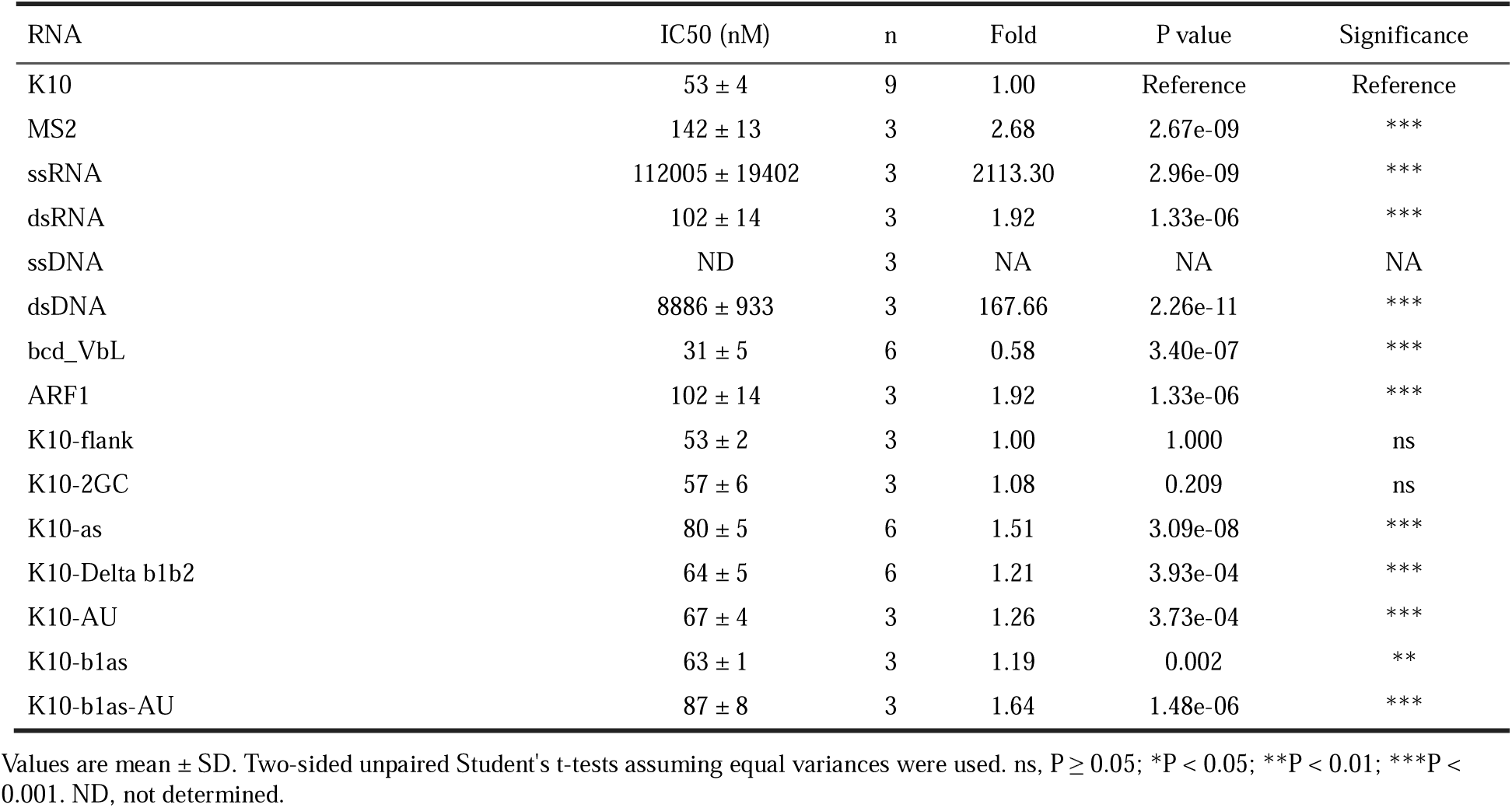
K10 versus other RNAs: IC50.

